# Freeze dehydration vs. supercooling in tree stems: physical and physiological modelling

**DOI:** 10.1101/2023.06.26.546491

**Authors:** Cyril Bozonnet, Marc Saudreau, Eric Badel, Thierry Améglio, Guillaume Charrier

## Abstract

Frost resistance is the major factor affecting the distribution of plant species at high latitude and elevation. The main effects of freeze-thaw cycles are damage to living cells and formation of gas embolism in tree xylem vessels. Lethal intracellular freezing can be prevented in living cells by two mechanisms: dehydration and deep supercooling. We developed a multiphysics numerical model coupling water flow, heat transfer, and phase change, considering different cell types in plant tissues, to study the dynamics and extent of cell dehydration, xylem pressure changes, and stem diameter changes in response to freezing and thawing. Results were validated using experimental data for stem diameter changes of walnut trees. The effect of cell mechanical properties was found to be negligible as long as the intracellular tension developed during dehydration was sufficiently low compared to the ice induced cryostatic suction. The model was finally used to explore the coupled effects of relevant physiological parameters (initial water and sugar content) and environmental conditions (air temperature variations) on the dynamics and extent of dehydration. It revealed configurations where cell dehydration could be sufficient to protect cells from intracellular freezing, and situations where supercooling was necessary. This model, freely available with this paper, could easily be extended to explore different anatomical structures, different species and more complex physical processes.

## Introduction

Plant species distribution at high latitude and elevation is mainly driven by their resistance to freezing stress (Charrier et al., 2013a). Freeze-thaw cycles are known to damage living cells (Siminovitch and Scarth, 1938; Mazur, 1969; Burke et al., 1976; Arora, 2018), as well as trees hydraulic functionning (Cochard et al., 1992; Utsumi et al., 1998; Cochard et al., 2001; Améglio et al., 2002; Charra-Vaskou et al., 2016). In this work, we focus on the former effect. Freezing stress induces damage in living cells by two main mechanisms: lethal intracellular freezing or frost-induced dessication (through cell membrane injury or protein denaturation), see Burke et al. (1976); Arora (2018); Pearce (2001). Living cells can prevent lethal intracellular freezing by two mechanisms: dehydration and deep supercooling (Levitt et al., 1980; Stegner et al., 2022).

Freeze-induced dehydration is induced by extracellular ice formation: upon decreasing temperatures, ice can form first in the extracellular medium and the low ice potential draws out cell water to growing extracellular ice masses (Olien, 1967; Sakai and Larcher, 2012). If dehydration is fast enough, intracellular sap concentration increases in living cells and lowers the phase change (freezing/melting) temperature, so that the living cell is protected against intracellular freezing (Towill and Mazur, 1976). Anti-freeze proteins can slightly lower the phase change temperature (Griffith and Yaish, 2004). Note however, as said previously, that prolonged dehydration itself can be a source of damage. A prerequisite for this strategy is that extracellular ice can form and grow. Some plants have evolved in order to keep space for extracellular ice to grow: specific expansion zones are found in leaves (*Eucalyptus pauciflora*, Ball et al. (2004)), between leaves inside buds (*Alnus alnobetula*, Neuner et al. (2019)), between bud scales (*Cornus officinalis*, Ishikawa and Sakai (1985)), in vallecular canals and pith cavity (*Equisitum hyemale*, Schott et al. (2017)), in bark (*Betula sp*, Schott and Roth-Nebelsick (2018)), or in xylem vessels (Utsumi et al., 1998; Neuner et al., 2010). How water may be directed towards these sites of ice accumulation may be linked with gradients of cell mechanical properties (Konrad et al., 2019). Acclimation of the plant to freezing stress also influences the formation and location of ice (Améglio et al., 2001a; Ball et al., 2004; Schott and Roth-Nebelsick, 2018). Increasing cooling rate, i.e., increasing the rate at which the external temperature is decreased, deteriorates the ability of the cell to dehydrate fast enough and to protect (by solute concentration) against intracellular freezing. For plant cells, 3°C/h is a sufficiently low cooling rate to ensure cell dehydration due to extracellular freezing (Arora, 2018).

If cellular dehydration is too slow, or does not happen, the temperature of intracellular sap may decrease below its melting point. Intracellular ice does not appear instantly, as a first ice crystal must form before ice to fill the entire cell. This state is known as supercooling, which becomes deep supercooling when the temperature of the sap is much lower than its melting point (difference of at least 10*−*°*C*, Cavender-Bares (2005)). Classical nucleation theory predicts that the probability for the first ice crystal to form increases with the volume of water in the compartment, the time spent in a supercooled state, a decrease in temperature, the presence of nucleation points such as heterogeneities, and other physical and geometrical factors, see Toner et al. (1990); Meng and Zhang (2020). Supercooling is observed in the extracellular medium as well (Améglio et al., 2001a; Neuner et al., 2010; Charra-Vaskou et al., 2016). Nucleation of supercooled water results in an exotherm (tracked by differential thermal analysis) (Burke et al., 1976; Neuner et al., 2010). Thus, when temperature decreases, a first exotherm related to extracellular freezing occurs, possibly followed by a second at much lower temperature, related to intracellular freezing (Burke et al., 1976). Intracellular ice nucleation temperature evolves seasonally and can be related to water and sugar contents (Charrier et al., 2013b, 2018; Baffoin et al., 2021), which are also related to management practices (Charrier et al., 2015).

Cell wall properties and geometry can influence the extent of freeze dehydration (Anderson et al., 1983; Bartolo et al., 1987; Wisniewski, 1995; Sakai and Larcher, 2012). Cell wall pores could also serve as an ice barrier (Ashworth and Abeles, 1984), thus preventing ice to spread in intracellular spaces. It is thought that negative turgor pressure, i.e., cell wall tension, generation can be linked to these properties, and limit the extent of dehydration, thus promoting supercooling (Rajashekar and Burke, 1996; Stegner et al., 2022). Several studies have directly linked cell wall properties to cold acclimation (Griffith and Brown, 1982; Stefanowska et al., 2002; Takahashi et al., 2021). Stegner et al. (2022) present a measure of these different parameters, as well as a measure of the extent of freeze dehydration. Through principal components analysis, they show that the extent of freeze dehydration is correlated negatively with cell elasticity, cell wall thickness, and the squared cell wall thickness to cell size ratio. These parameters can indeed influence negative turgor pressure generation through their influence on cell collapse pressure (Ding et al., 2014).

Freeze-induced dehydration dynamic at the cellular scale can be observed at the tissue scale through changes in stem diameter (Améglio et al., 2001a; Zweifel and Häsler, 2000). They result from the migration of liquid water from elastic living cells to frozen rigid apoplastic tissues. This dehydration of elastic living cells results in volume changes at the tissue level, hence to diameter changes. The magnitude and reversibility of such changes is related to the degree of cold acclimation of the stem and to the temperature imposed (Améglio et al., 2001a). Diameter changes are mainly due to bark thickness changes, but the xylem tissue shrinks as well (Améglio et al., 2001a; Charra-Vaskou et al., 2016; Lintunen et al., 2017), in a much smaller proportion than bark tissue (the xylem shrinkage accounts for less than 30% of the total shrinkage).

Xylem pressure variations have been observed in stems in response to freezing and to freeze-thaw cycles (Améglio and Cruiziat, 1992), and are either believed to be due to ice volumetric expansion (Robson and Petty, 1987) or to flow between apoplastic elements (O’Malley and Milburn, 1983; Tyree, 1983; Milburn and O’Malley, 1984). These pressure variations may be due to the ideal gas law (Tyree, 1983), i.e., they may originate from the contraction or expansion of gas within the apoplast (probably in vessels) due to water fluxes and temperature changes. The level of hydration of the stem also influences these pressure variations (Milburn and O’Malley, 1984).

Several existing approaches separately modelled frost hardiness, diameter and xylem pressure changes in response to freezing stress. Frost hardiness was modelled using physiological parameters, such as water and sugar content (Poirier et al., 2010; Charrier et al., 2013b, 2018; Baffoin et al., 2021). It was also modelled using variables such as temperature, photoperiod and phase of annual development (Leinonen, 1996). Freeze-induced dehydration has been modelled in Rajashekar and Burke (1996), where the link between wall tension and living cell dehydration was studied, and more recently in Konrad et al. (2019), where cell water relations were examined in case of the presence of water in different phases (liquid, vapour, solid) and of a gradient of mechanical properties accross living cell tissues. In Eurich et al. (2021), a finite element porous model was developed to study cell dehydration and tissue shrinkage induced by extracellular freezing.

Diameter changes have been modelled for fully frozen stems by using a two compartment model (xylem and bark) by Lindfors et al. (2019). In the latter reference, the xylem potential was equal to the ice potential, and any change in temperature resulted in change in ice potential that was matched by the bark potential (sum of turgor and osmotic potential), thus inducing diameter changes. By fitting numerical results onto experimental data, the authors found a variation of cell wall elasticity with temperature. The magnitude of living cell turgor pressure was not shown, but as turgor loss was not accounted for in the model, cell wall tensions were likely to be generated. We can thus safely assume that the strategy predicted by their model for intracellular freezing avoidance is supercooling, which was not confirmed, nor refuted, by experimental results.

Xylem pressure changes have been modelled for a maple stem subjected to freeze-thaw cycles (Ceseri and Stockie, 2013; Graf et al., 2015). In the latter reference, phase change and other processes like gas and water transport at the microscopic scale were coupled to heat transport at the macroscopic scale, and successfully reproduced the pressure changes observed in O’Malley and Milburn (1983). This model was furthermore coupled to an asymmetric root uptake and reproduced the pressure build-up over multiple freeze-thaw cycles observed in Ewers et al. (2001); Améglio et al. (2001b). The work in Ceseri and Stockie (2013); Graf et al. (2015) did not include living cells.

In the present work, we study the dynamic and extent of living cell dehydration induced by extracellular freezing, along with its coupling with xylem pressure and stem diameter changes. The ultimate goal is to enhance our understanding of the competition between freeze dehydration and supercooling for the protection against intracellular freezing. To accomplish this, we used a modelling approach.

We have developed a comprehensive physical model that integrates water and heat fluxes, extracellular phase change, and living cell dehydration across various tissues. Additionally, the model incorporates living cell pressure-volume relationships, including the impact of turgor loss, as well as changes in stem diameter.

For validation purpose, we compared the outputs of the model to experimental data for diameter changes of stems exposed to freezing stress. Then, we investigated the influence of mechanical properties, environmental conditions, and physiological variables on the dynamics and extent of cell dehydration, and on pressure and diameter changes.

Finally, we presented a thorough analysis of the model results and discussed them in the context of existing literature, highlighting key findings and potential avenues for further research. The model is freely available along with the paper so that others could contribute to its development and benefit from its use.

## Numerical model

### General description

The model relies on a wood anatomy description in the transversal plane of a wood section. The anatomy of a walnut branch is described by Alves (Alves et al., 2001, 2007) and is presented in figure 1, at the tissue (figure 1a), and cell (figure 1b) scales. The structure of our model (figure 1c) groups the essential anatomical elements to simulate the processes described above: xylem vessels, vessel-associated cells (VACs) and bark cells. It shares some similarities with the model of Hölttä et al. (2006): living cells are interconnected by a parenchyma ray, connected at its periphery to the bark cells, and connected to a radial alignment of vessels. We assume that the external temperature field is homogeneous around the stem, i.e. axi-symmetric, so that radial exchanges are only modelled along one ray and multiplied at the xylem/bark interface by *N*_*ray*_, the number of parenchyma rays. The longitudinal dimension is not considered. The parenchyma ray itself is not described explicitly, i.e., individual (isolated) ray cells are omitted, but rather represented by a hydraulic resistance between VACs, and between VACs and bark cells. Water flows and volume changes are computed for one VAC per vessel and for one bark cell in the bark tissue. These water flows and volume changes are then rescaled by *N*_*vac*_, the number of VACs connected to each vessel, and *N*_*Bark cell*_, the number of living cells in the bark, similarly to what is done in Graf et al. (2015) for the fiber/vessel fluxes.

**Figure 1:**
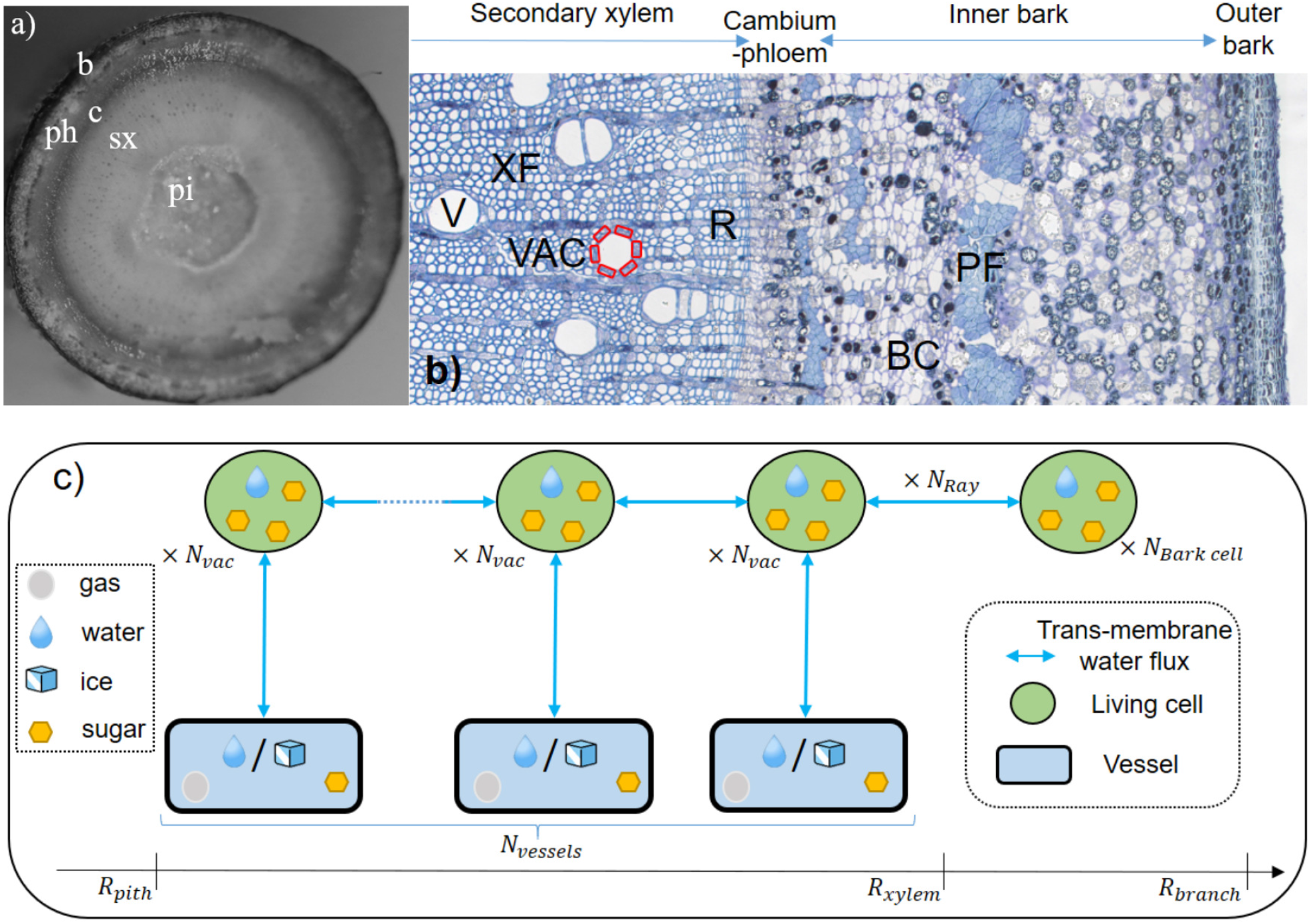
a) *Juglans regia* branch anatomy at the tissue scale. sx: secondary xylem, pi: pith tissue, c: cambium, ph: phloem, b: bark. b) *Juglans regia* branch anatomy at the cellular scale. XF: xylem fibers, V: vessel, R: parenchyma ray, BC: bark cell, PF: phloem fibers, VAC: vessel associated cells (highlited in red). c) Model structure. Depending on the type of element and its enthalpy level, water is assumed to be in different phases (solid, liquid, gas). Water fluxes (blue arrows) occur between different anatomical elements across cell membranes. Scaling coefficients (*N*_*vac*_, *N*_*Ray*_, *N*_*bark cell*_, *N*_*vessels*_) are used to obtain a more accurate anatomical description.

Elastic living cells, i.e., bark cells and VACs, are assumed to contain only water and soluble sugar. We therefore assume that intracellular ice does not form in the temperature range we study, which is discussed a posteriori depending on the level of dehydration and/or supercooling degree. Rigid vessels contain sugar, liquid water or ice depending on local temperature, and gas. This gas compresses or expands in response to water flows entering or leaving the vessels, thus creating pressure variation according to the ideal gas law, as done in Ceseri and Stockie (2013); Graf et al. (2015).

Heat transfer and phase changes are calculated at the tissue scale and driven by external temperature variations. Temperature (and phase for xylem sap) is computed at the cellular scale from the tissue-scale values. Vessel sugar content impacts tissue-scale phase change through freezing point depression (FPD).

Water fluxes occur between the different elements, see the blue arrows in figure 1c. These water fluxes are determined by the differences in water potential (hydrostatic/turgor, osmotic, cryostatic) across cell membranes. For each elastic compartment (VACs, bark cells), the balance of water fluxes results in volume changes, which are then used to calculate changes in tissue diameter, as well as changes in turgor and osmotic potential.

## Mathematical description

### Anatomy

The anatomical description used in the model is shown in figure 1c. Vessels are arranged regularly along the ray, with the vessel number, *N*_*vessels*_, computed using a linear vessel density and the size of the xylem tissue:

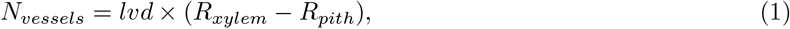

where *lvd, R*_*xylem*_ and *R*_*pith*_ are the linear vessel density, the xylem and the pith radius, respectively. Each vessel has a given number of VACs associated with it, *N*_*vac*_, calculated using a ratio of the vessel-VAC exchange area to the projected VAC area:

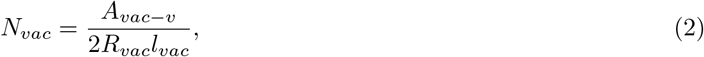

where *A*_*vac−v*_, *R*_*vac*_ and *l*_*vac*_ are the vessel-VAC exchange area, the VAC radius and VAC length, respectively. The number of parenchyma rays is computed using a tangential ray density and the branch diameter:

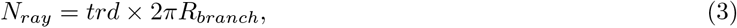

where *trd* is the tangential ray density. The number of bark cells connected to the parenchyma rays is

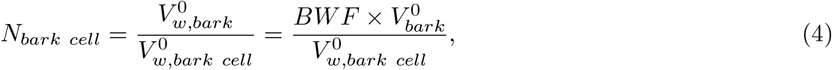

with 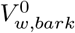 the inital volume of water in the bark accesible from the rays, equal to 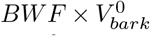, with *BWF* the bark water fraction accessible from the rays, and 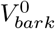 the initial bark volume. 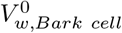 is the initialbark cell water volume.

### Heat transfer and phase change

Heat transfer and phase change are calculated at the tissue scale through the heat equation in a 1D axi-symmetric model in cylindrical coordinates:

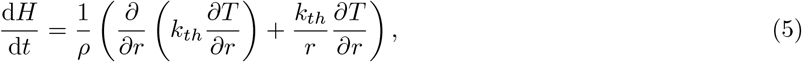

where H is the enthalpy, T the temperature, *k*_*th*_ the thermal conductivity and *ρ* the density. This equation is used for *r* in]*R*_*pith*_; *R*_*xylem*_[and completed with the following boundary conditions (Graf et al., 2015):

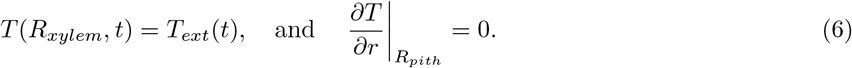

Equation (5) must be completed by thermodynamic relationship between *H* and *T* in order to account for phase change. This is done by using regularized *H*-*T* relationship, as done in Graf et al. (2015) and shown in figure 2a, where the jump in enthalpy at the melting point corresponding to latent heat release or storage is spread over a very small temperature interval (in our case Δ*T*_*m*_ = 0.003K) for numerical convenience. Density and thermal conductivity are also computed locally based on the enthalpy value, see figure 2b. The phase change temperature at the tissue scale is computed locally based on 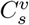, the local vessel sugar content:

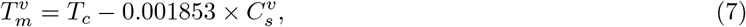

with *T*_*c*_ = 273.15K. Note that Eq. (7) is only valid for water. Heat equation, Eq. (5), is solved on a grid finer than that formed by the regular arrangement of vessels along the ray, but with grid points that correspond to the vessel locations, so that temperature and enthalpy values do not need to be interpolated to be obtained at the cell scale. The cell size of the thermal grid, Δ_*r*_, is small enough to reach spatial convergence of the results. The vessel sugar concentration is linearly extrapolated from the vessel locations to the heat equation grid so that Eq. (7) can be applied at any thermal grid point. A vessel ice fraction, *δ*_*iv*_, is computed for each vessel as a function of the local enthalpy value, as shown in figure 2b.

**Figure 2:**
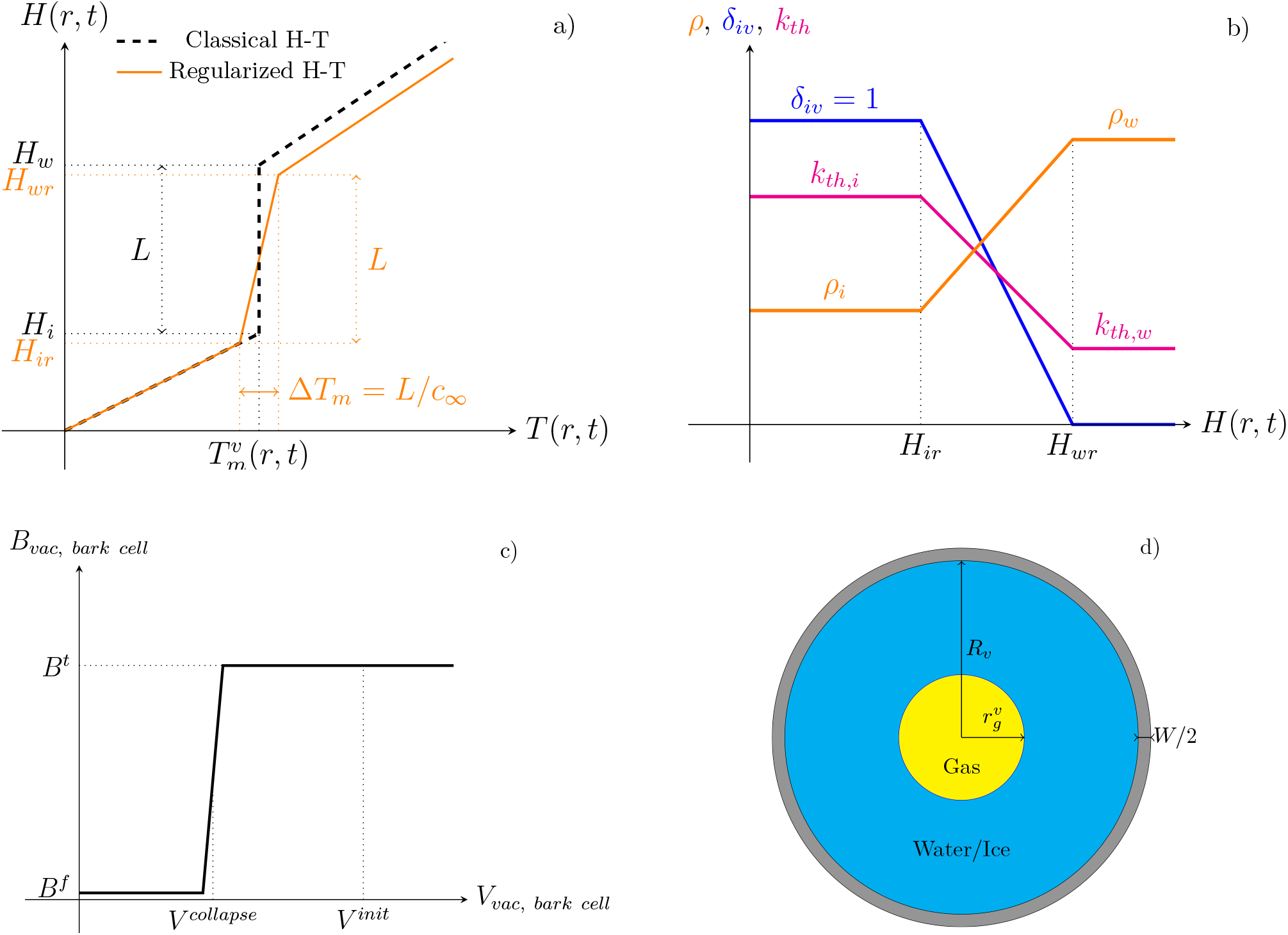
a) Enthalpy-Temperature relationship: the regularized relationship spreads latent heat release/storage over a very small temperature interval of thickness Δ*T*_*m*_ = *L/c*_∞_ . In the ice or liquid state the enthalpy is a linear function of temperature, i.e., *H* = *c*_*i*_*T* or *H* = *c*_*w*_(*T −*(*T*_*m*_ + Δ*T*_*m*_*/*2)) + *L*, with *c*_*i*_ and *c*_*w*_ the ice and liquid water specific heat capacities, respectively. b) Changes of physical properties and vessel ice fraction with local enthalpy value. c) Changes of cell elastic modulus with cell volume, from *B*^*t*^ the value for a turgid cell, to *B*^*f*^ the value for a flacid cell. Around *V*_*collapse*_, the elastic modulus is regularized in the same way as the H-T relationship around the melting temperature. d) Geometry of a vessel of radius *R*_*v*_, wall thickness *W/*2, containing a gas bubble of radius 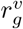.

### Water fluxes

Water fluxes between elements are computed using Darcy’s law. Along the parenchyma ray:

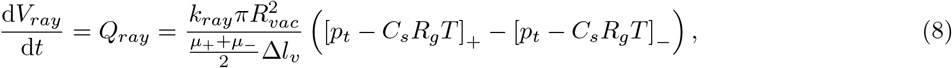

where *k*_*ray*_ is the ray porosity, Δ*l*_*v*_ is the distance along the ray between two vessels, *µ* is the dynamic viscosity of the water and sugar solution computed locally using the law given in Chenlo et al. (2002), *p*_*t*_, *C*_*s*_ and *T* are the living cell (VAC or bark cell) turgor pressure, sugar concentration and temperature, respectively. The + and*−*signs represent a differentiation along the ray from the inside to the oustide of the stem (up to the bark cells). *Q*_*ray*_ is positive for water fluxes going towards the inside of the branch. Between one vessel and one corresponding VAC, the flow rate is

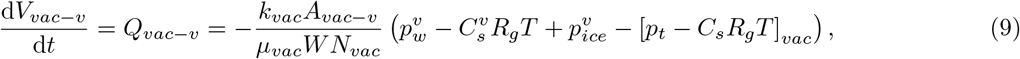

where *k*_*vac*_ is the vessel-VAC wall porosity, *W* the vessel-VAC wall thickness, and 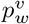 the vessel water pressure.*Q*_*vac*_*−*_*v*_ is positive for water fluxes going towards the vessels. The cryo-suction pressure induced by vessel freezing is computed at each vessel location as (Loch, 1978; Beck et al., 1984)

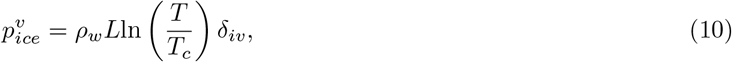

where *ρ*_*w*_ and *L* are the water density, and water latent heat of fusion, respectively.

Eq. (9) implies that cryo-suction will draw water in a vessel from its VACs once this vessel is frozen. It is assumed that this water turns into ice when entering a vessel. We furthermore assume that, if a vessel is frozen, i.e., if *δ*_*iv*_ = 1, *Q*_*vac−v*_ cannot be negative.

### Pressure-volume relationships in living cells

In living cells, the balance of water fluxes results in volume changes. For the VACs:

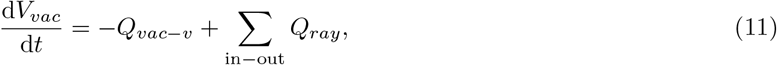

where the second term on the right hand side represents the balance of fluxes entering/leaving the VAC from/to the ray. Between the xylem tisue and the bark, the total water flux is

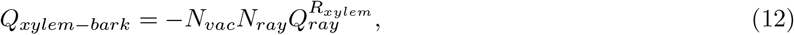

where 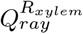 is the water flux computed between one bark cell and the VAC closest to the bark. This water flux is rescaled by the number of bark cells to obtain the volume change at the bark cell scale:

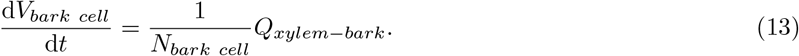

These living cells volume changes are related to turgor pressure variation through (Steudle et al., 1977)

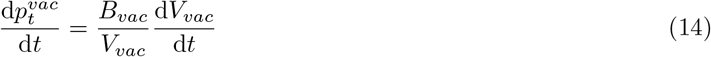

for the VACs, and

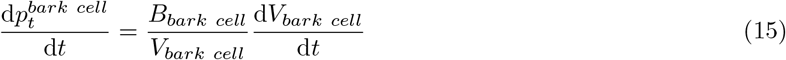

for the bark cell. In the previous two equations, *B*_*vac*_ and *B*_*bark cell*_are the VAC and bark cell elastic modulus, respectively. Cell turgor loss, i.e., cell collapse in case of reversible cytorrhysis (Arora, 2018), occurs either for *p*_*t*_ = 0, or at a negative turgor pressure value whose magnitude depends on cell dimension and mechanical properties (Ding et al., 2014; Yang et al., 2017). To accurately model cell turgor loss and recovery we choose to impose a relationship between cell elastic modulus and cell volume, as shown in figure 2c. Particularly, the elastic modulus drops to a very low value, *B*_*f*_, when the cell reaches the volume at which it collapses. This volume is computed as

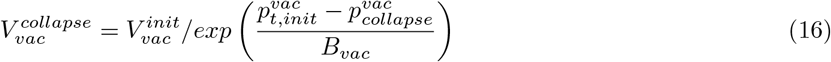

for the VACs, where 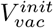 is the initial water volume in the cell, 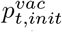 the initial turgor presure and 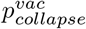 the pressure at which the cell collapses, respectively. A similar equation is employed for the bark cell. Eq. (16) comes from the integration of Eq. (14) between the initial state and cell collapse. Above this critical volume, for simplicity and for the validity of Eq. (16), the elastic modulus is assumed to be constant, even though experimental results suggest it could vary with turgor pressure and/or cell volume (Steudle et al., 1977).

Volume changes also result in osmotic pressure changes through changes in sugar concentration:

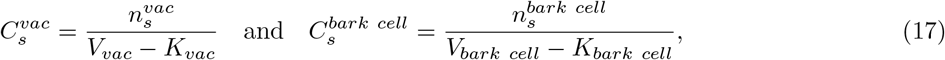

where 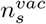and 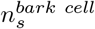 are the sugar quantity in the VAC and bark cell, and *K*_*vac*_, *K*_*bark cell*_, are the cell volume where no sugar can be contained (certain cell organelles), respectively. As no sugar fluxes are considered in the present work, the sugar quantities are constant. These changes in sugar concentration result in living cell FPD for the VACs,

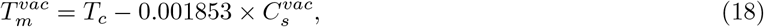

and for the bark cells,

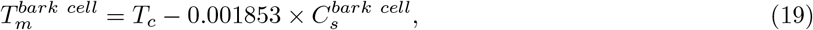

with 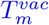, the freezing temperature computed for each VAC, and 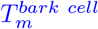, the freezing temperature for the bark cells. Note again that these last two equations are only valid for water.

### Pressure-volume relationships in vessels

Vessels contain gas that compresses or expands depending on water fluxes leaving or entering vessels. Following Ceseri and Stockie (2013); Graf et al. (2015), and as shown in figure 2d, we assume that this gas is contained in one cylindrical bubble located at the center of each vessel. Applying flow rate conservation between the gas/water (or gas/ice) interface and the vessel/VAC wall, the bubble radius, 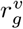, varies as

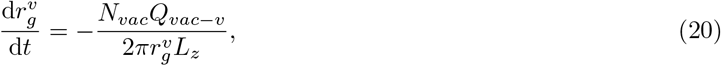

where *L*_*z*_ is a vertical dimension that is introduced for unit consistency but that has no influence on model results. Changes in 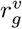induce changes in gas pressure through the ideal gas law:

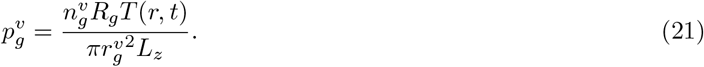

In previous equation, 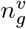 is the gas quantity inside the gas bubble and *R*_*g*_ the ideal gas constant. Finally, the pressure in the liquid water/ice phase is obtained using Laplace equation:

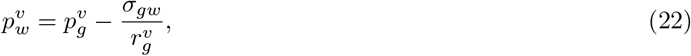

where *σ*_*gw*_ is the liquid water/gas interface surface tension. Note that we assume that no change occurs in Eq.(22) when the vessel is entirely frozen. Similarly to living cells, volume changes in vessels also result in sugar concentration changes:

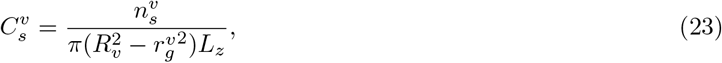

where 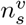 and *R*_*v*_ are the vessel sugar quantity, taken as a constant, and vessel radius, respectively. These changes in vessel sugar concentration impact phase change at the tissue scale through Eq. (7).

### Diameter changes

Finally, diameter changes are obtained from living cell volume changes, Eqs. (11) and (13). The total volume of water in VACs is computed at each instant:

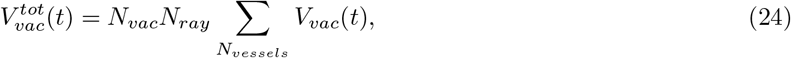

with the volume variation equals to

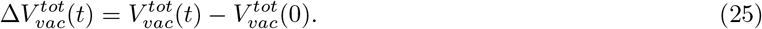

The xylem diameter, considered as a cylinder, is computed as

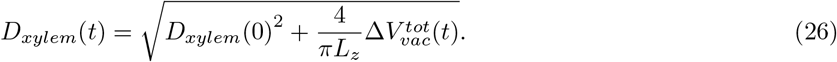

At each instant, the volume of the bark tissue is equal to the initial volume minus the volume lost by dehydration:

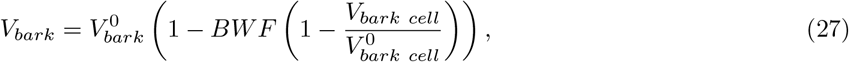

from which the stem diameter, considered as a cylinder, can be computed:

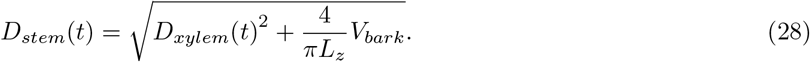

### Numerical resolution and parameter choices

The model is implemented in the Matlab software version R2018a (MATLAB, 2018). Spatial discretisation of Eq. (5) is ensured using the finite difference method. The system of differential equations formed by equations (5), (8), (9), (11), (13), (14), (15) and (20) is advanced in time using Matlab’s variable order *ode*15*s* solver based on numerical differentiation formulas, which is specifically designed for stiff equations, with a maximal time step of 1*s*, which is a sufficient value to resolve any stiffness in the problem under study. The other equations are state equations computed at each time step. Note that we verified the implementation of Eq. (5) using an analytical solution (Prapainop and Maneeratana, 2004) for a 1D freezing-front propagation (Stefan problem, R^2^ *>* 0.9999), and using a reference finite element solver (Comsol Multiphysics (COMSOL, 2020)) for the 1D axi-symmetric implementation (R^2^ = 0.9998). The reference result presented in figure 3 takes a computational time of around 5 minutes on a Dell Latitude 7490 with 1.7 GHz quad-core Intel i5 processor. The source code can be downloaded at https://github.com/cyrilbz/freezing_branch.

**Figure 3:**
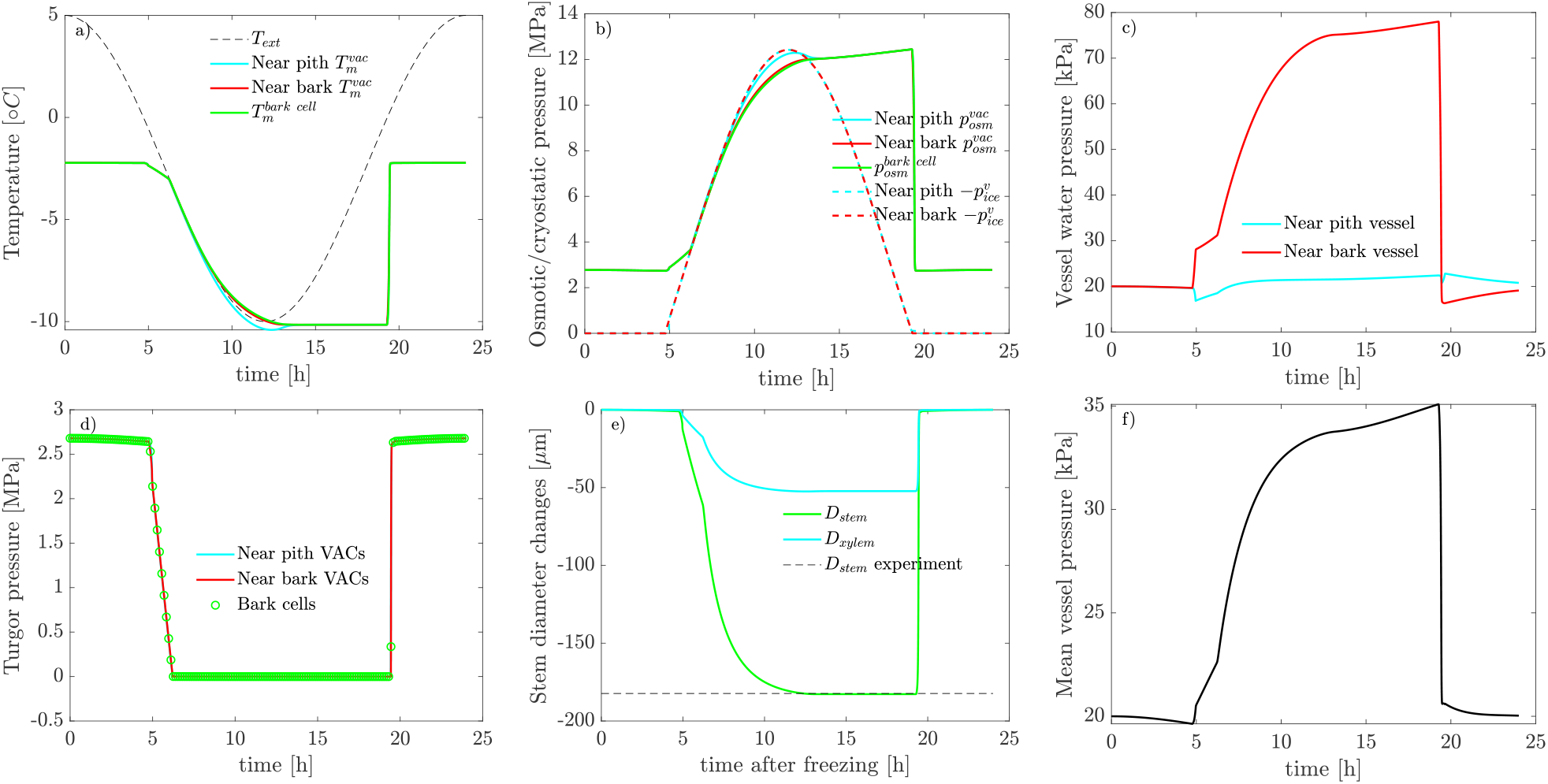
Model results for one freeze-thaw cycle. The stem (2 cm in radius), initially at +5°C, is subjected to a sinusoidal external temperature variation, down to *−*10°C, over 24 hours. The parameters used for this case correspond to base case 1 in table 2. a) FPD dynamic for two VACs and one bark cell, along with external temperature changes. b) Time course of osmotic pressure for two VACs and one bark cell, and cryostatic pressure for two vessels. c) Time course of 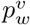 for two vessels. d) Time course of living cell turgor pressure for two VACs and one bark cell. Note that for the bark cell, only one data point every 50 is shown to enhance readability. e) Tissue diameter changes (total stem and xylem) and value given by the correlation in (Améglio et al., 2001b). f) Mean vessel pressure.

All model and state variables are regrouped in table 1. All parameters are in table 2. All values are either justified based on the literature, or have been specifically measured or calibrated, except the initial sugar content in living cells, *k*_*ray*_, *B*_*f*_, 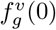, and the thermal conductivities. The initial sugar content in living cells has been estimated based on measurements in Charrier et al. (2013b). The ray hydraulic conductivity, *k*_*ray*_, has been chosen high enough so that water volumes coming from the bark are more evenly distributed across the xylem tissue and not located in the vicinity of the bark only. The flacid elastic modulus, *B*_*f*_, has no impact on the simulation results as long as it is small in front of *B*_*t*_. The vessel gas volume fraction, 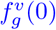, has been chosen so that vessel pressure variations have an order of magnitude consistent with the experiments of Améglio et al. (2001b). Finally, the thermal conductivities are chosen one order of magnitude above their physical values so that the ice front propagation speed is more realistic (higher than with the physical values) and the repartition of the water volume coming from the living cells is more homogeneous across the vessels. Increasing the ice front speed reduces the time between the freezing of two successive vessels, hence at a given time it can increase the number of locations for ice accumulation.

**Table 1:**
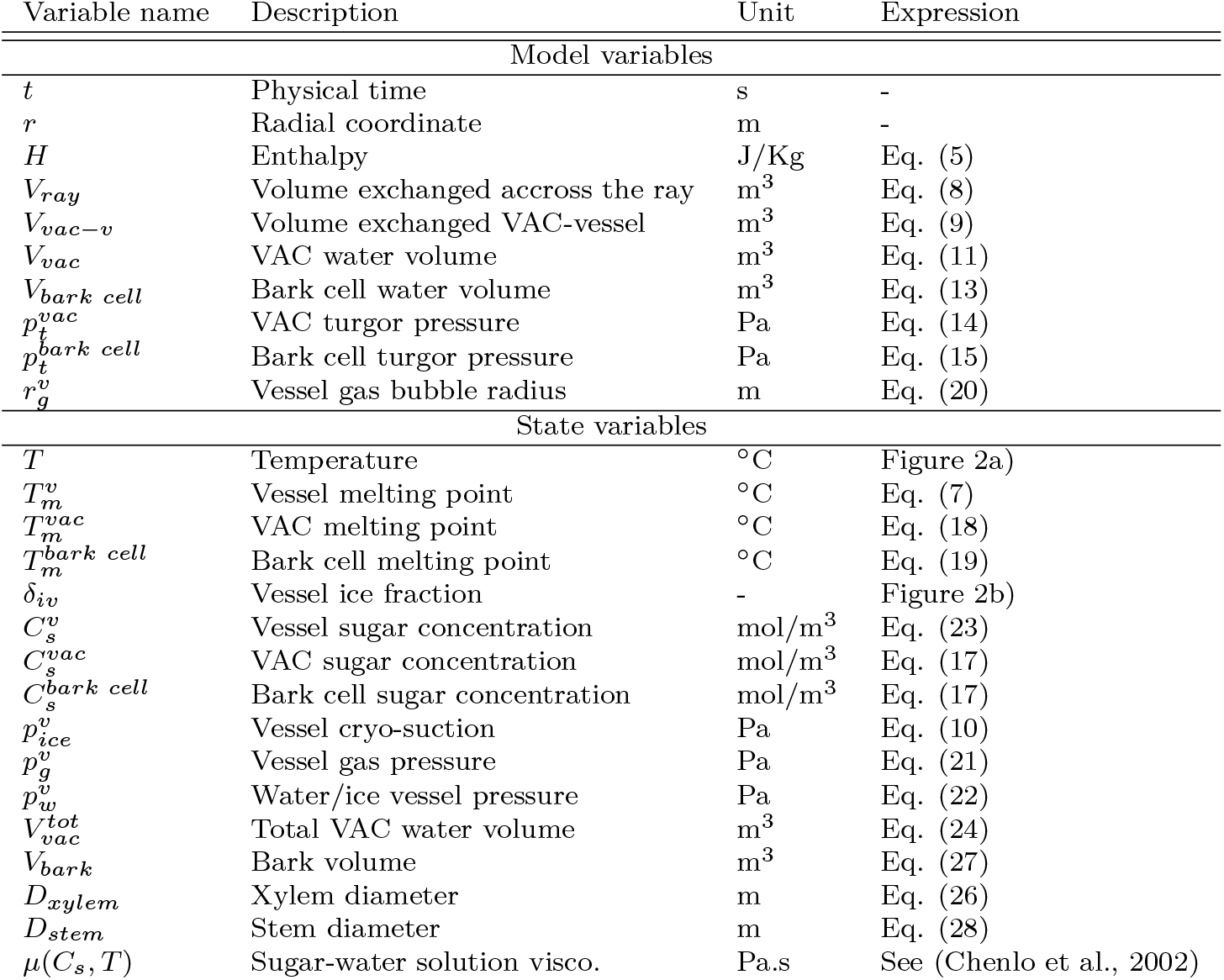
Model and state variables.

| Variable name | Description | Unit | Expression |
| --- | --- | --- | --- |
| Model variables |  |  |  |
| $t$ | Physical time | s | - |
| $r$ | Radial coordinate | m | - |
| $H$ | Enthalpy | J/Kg | Eq. (5) |
| $V_{ray}$ | Volume exchanged accross the ray | m<sup>3</sup> | Eq. (8) |
| $V_{vac-v}$ | Volume exchanged VAC-vessel | m<sup>3</sup> | Eq. (9) |
| $V_{vac}$ | VAC water volume | m<sup>3</sup> | Eq. (11) |
| $V_{bark\ cell}$ | Bark cell water volume | m<sup>3</sup> | Eq. (13) |
| $p_t^{vac}$ | VAC turgor pressure | Pa | Eq. (14) |
| $p_t^{bark\ cell}$ | Bark cell turgor pressure | Pa | Eq. (15) |
| $r_g^v$ | Vessel gas bubble radius | m | Eq. (20) |
| State variables |  |  |  |
| $T$ | Temperature | °C | Figure 2a) |
| $T_m^v$ | Vessel melting point | °C | Eq. (7) |
| $T_m^{vac}$ | VAC melting point | °C | Eq. (18) |
| $T_m^{bark\ cell}$ | Bark cell melting point | °C | Eq. (19) |
| $\delta_{iv}$ | Vessel ice fraction | - | Figure 2b) |
| $C_s^v$ | Vessel sugar concentration | mol/m<sup>3</sup> | Eq. (23) |
| $C_s^{vac}$ | VAC sugar concentration | mol/m<sup>3</sup> | Eq. (17) |
| $C_s^{bark\ cell}$ | Bark cell sugar concentration | mol/m<sup>3</sup> | Eq. (17) |
| $p_{ice}^v$ | Vessel cryo-suction | Pa | Eq. (10) |
| $p_g^v$ | Vessel gas pressure | Pa | Eq. (21) |
| $p_w^v$ | Water/ice vessel pressure | Pa | Eq. (22) |
| $V_{vac}^{tot}$ | Total VAC water volume | m<sup>3</sup> | Eq. (24) |
| $V_{bark}$ | Bark volume | m<sup>3</sup> | Eq. (27) |
| $D_{xylem}$ | Xylem diameter | m | Eq. (26) |
| $D_{stem}$ | Stem diameter | m | Eq. (28) |
| $\mu(C_s, T)$ | Sugar-water solution visco. | Pa.s | See (Chenlo et al., 2002) |

**Table 2:**
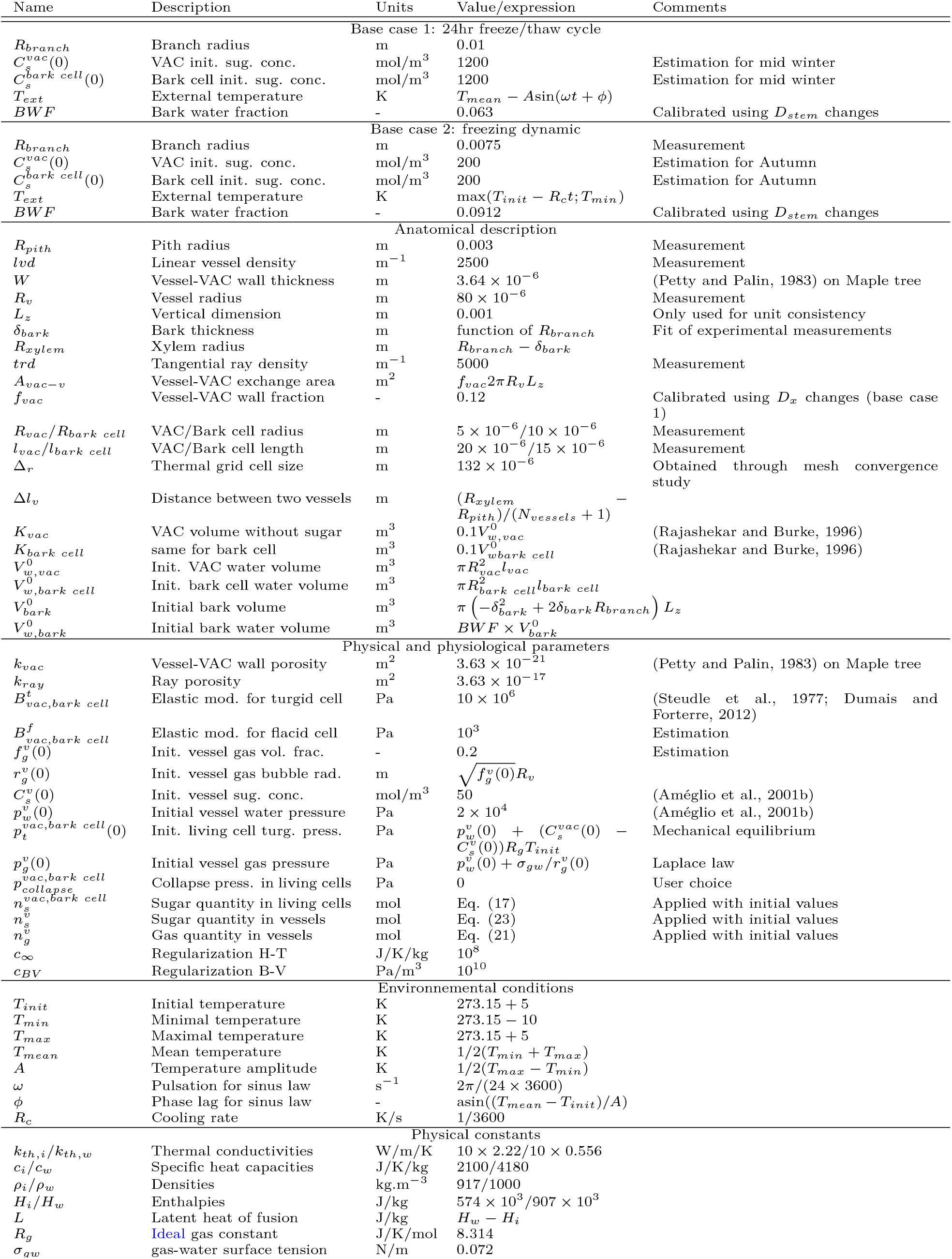
Parameter list, description & values. Measurements and estimations were done for *Juglans Regia* stems.

For simplicity, we further assume that all living cells have the same mechanical properties and the same initial sugar concentrations, but the model is already capable of handling different parameter values between VACs and bark cells.

## Results

In the present section we present the results obtained using the model described previously. Unless stated otherwise, all parameter values are presented in table 2.

### Freeze-thaw cycle

In the present section we introduce the model result for a 24 hours long freeze-thaw cycle (base case 1 in table 2). The case simulated corresponds to a walnut stem with a diameter of 2 cm, initially at +5°C and subjected to a sinusoidal external temperature variation, down to −10°C, over 24 hours. The external temperature profile is given in figure 3a. At *t* = 4.77h, freezing starts in the vessel closest to the bark, see the progressive increase in the cryosuction close to the bark 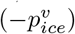 in figure 3b. It draws liquid water from adjacent cells to the vessels, which result in a pressure increase in the vessels, see figure 3c. Simultaneously, the liquid content of the vessel closest to the pith is attracted towards the ice front. Sap thus flows out of the vessel through the rays towards frozen vessels, which induces a slight pressure decrease (figure 3c). At *t* = 4.96h, the vessel closest to the pith freezes and the whole xylem tissue is frozen. Phase change is completed and the only remaining thermal phenomenon is diffusion, which in a few seconds brings all vessels to equilibrium in temperature and cryostatic pressure (figure 3b). After this step, for the vessel closest to the pith the pressure increases too, as water is entering the vessel due to cryostatic suction (figure 3c).

As freezing starts, the turgor pressure in living cells decreases (figure 3d) and their osmotic potential increases (figure 3b). Whatever their positions, all living cells have a very close dehydration dynamic resulting in only minute differences in turgor pressure between VACs (close to the bark or the pith) and bark cells. The loss of turgor occurs around *t* = 6.25h and living cells collapse (reversible cytorrhysis). It can be seen on all variables presented in figure 3 as a tipping point in dehydration dynamic: as soon as turgor loss occurs, living cell dehydration becomes faster, and the osmotic potential of living cells nearly matches the vessel cryostatic pressure (figure 3b).

The dehydration of living cells lowers the freezing point depression (FPD), as seen in figure 3a. The initial FPD value depends on the initial living cell sugar content. FPD dynamic tightly follows external temperature variation, and dehydration is fast enough to limit intracellular freezing probability, i.e., melting temperature based on living cell sugar concentration is almost always lower than the external temperature.

Cell dehydration from freezing also induces tissue diameter shrinkage, see figure 3e. Turgor loss point is observed when shrinkage accelerates (around *t* = 6.25h). Beyond this point, shrinkage becomes progressively slower and stabilizes around 180*µ*m, i.e., the value given by the correlation in Améglio et al. (2001a). Bark tissue shrinkage is largely dependant on *BWF*, the bark water fraction accessible from the rays, which has been calibrated to obtain the desired shrinkage. Similarly, xylem tissue shrinkage largely depends upon *f*_*vac*_, the fraction of the vessel wall covered by VACs, which ultimately controls the amount of water that can move from elastic cells (VACs) to rigid elements (vessels). *f*_*vac*_ has thus been calibrated in order to obtain the target xylem shrinkage, i.e., 29% of the total stem shrinkage according to the correlation given in Améglio et al. (2001a).

Once temperature increases, cryostatic suction starts decreasing (incrasing ice potential) but as we hypothetized that ice does not flow out of vessels, the osmotic potential diverges from the decrease in cryosuction. It does rather slightly increase with temperature, as vessel water pressure does, in relation to temperature effect in ideal gas law. As the dehydration of living cells ceased, tissue diameter and FPD remained stable until the temperature reaches the vessel melting point.

When thawing starts, the difference in osmotic potential between living cells and vessels is so high that the swelling of the diameter is extremely rapid. As no irreversible process takes place, e.g., cell mortality due to intracellular freezing, diameter and pressure changes are fully reversible.

The mean vessel pressure signal is shown in figure 3f. It progressively increases after freezing start, reaching values up to 35kPa, which is an order of magnitude similar to that in Améglio et al. (2001b). It then drops when thawing starts.

### Validation against experimental results

In this section we validate the model against experimental stem diameter changes of *Juglans Regia*. The first test corresponds to a verification of the skrinkage dynamic after freezing inception. The experimental result for shrinkage dynamic has been obtained with the method described in Améglio et al. (2001a) and at the same period (November). In figure 4a, one can see that except at short term, modelled stem shrinkage clearly follows the experimental results. Similarly to what has been described in previous section, turgor loss occurs around *t−t*_*freezing*_ = 0.28h, characterized by a change in the rate of shrinkage.

**Figure 4:**
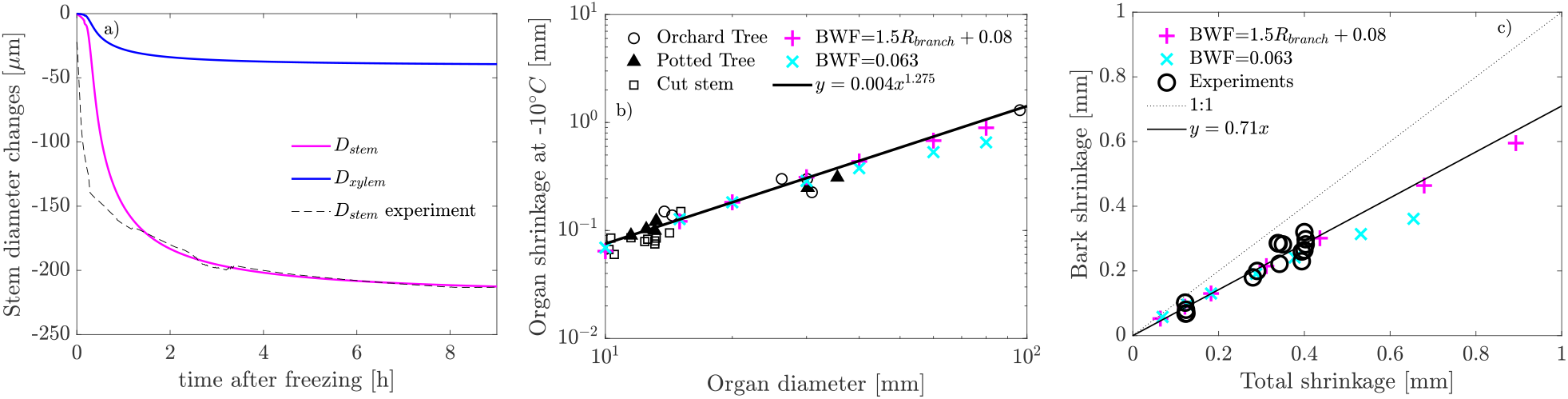
Validation of the model against experimental results. a) Shrinkage dynamic after freezing. The parameters used for this case correspond to base case 2 in table 2. The initial diameter is the diameter measured at room temperature before the experiment. b) Effect of organ diameter (2*R*_*branch*_) on maximal shrinkage at *−*10°C. c) Bark versus total shrinkage. In b and c we compare between the model results for two BWF modelling (constant or variable with branch radius) with experimental results and correlations from (Améglio et al., 2001a), for the parameters corresponding to base case 1 in table 2.

As the model is performant at predicting middle and long term stem shrinkage dynamic we now focus at predicting the effect of organ diameter (2*R*_*branch*_) on maximum shrinkage at 10°C. As shown on figure 4b, the model accurately predicts the relation of maximal shrinkage with diameter, see also the R^2^ values in table 3. The error increases for thick branches, especially when *BWF* is constant. The increase in shrinkage with organ diameter is due to the increase in the amount of water accessible from the ray with diameter through the increase in *δ*_*bark*_ and, if not taken constant, *BWF* .

**Table 3:** Determination coefficient values between experimental correlations and experimental or model results.

| | $y = 0.004x^{1.275}$ (figure 4b) | $y = 0.71x$ (figure 4c) |
| --- | --- | --- |
| Experiments | $R^2 = 0.9809$ (n=25) | $R^2 = 0.8995$ (n=16) |
| Model $BWF = 0.063$ | $R^2 = 0.2196$ (n=7) | $R^2 = 0.7949$ (n=7) |
| Model $BWF = f(R_{branch})$ | $R^2 = 0.9392$ (n=7) | $R^2 = 0.9918$ (n=7) |

Finally, the ratio between bark and total shrinkage is studied figure 4c. The model is particularly good at predicting the difference of magnitude between total and bark shrinkage, particularly when *BWF* is not constant, see table 3 for R^2^ values. As *D*_*xylem*_ does not vary with *BWF*, it means that xylem diameter changes are computed correctly, whatever the choice for *BWF* . The error in stem diameter changes in figure 4b is mainly explained by the error in bark shrinkage.

Overall, the model is capable of predicting middle and long term shrinkage dynamic and amplitude (at *−*10°C) as well as the relative contributions of the different tissues, with only two calibrated parameters (*BWF*,*f*_*vac*_).

### Effect of mechanical properties

We now study the effect of mechanical parameters on the dehydration dynamic, xylem pressure and stem diameter changes. We start by describing the effect of the turgid elastic modulus, *B*_*t*_, i.e., the elastic modulus before cell collapse. As shown in figures 5a-b-c, the value of *B*_*t*_ does not have any impact on the middle and long term results on the living cell freezing point depression, the mean vessel pressure and the stem diameter. It does actually only have an effect before turgor loss, as shown with the insets in figures 5b and c. Lowering the turgid elastic modulus results in a faster stem shrinkage when the cell is still turgid, and a delayed cell collapse. These changes can also be seen in the mean vessel pressure dynamic, with a faster increase at short term for lower *B*_*t*_. Beyond turgor loss, no effects of *B*_*t*_ is noticeable on stem dehydration, freezing point depression, and mean vessel pressure.

**Figure 5:**
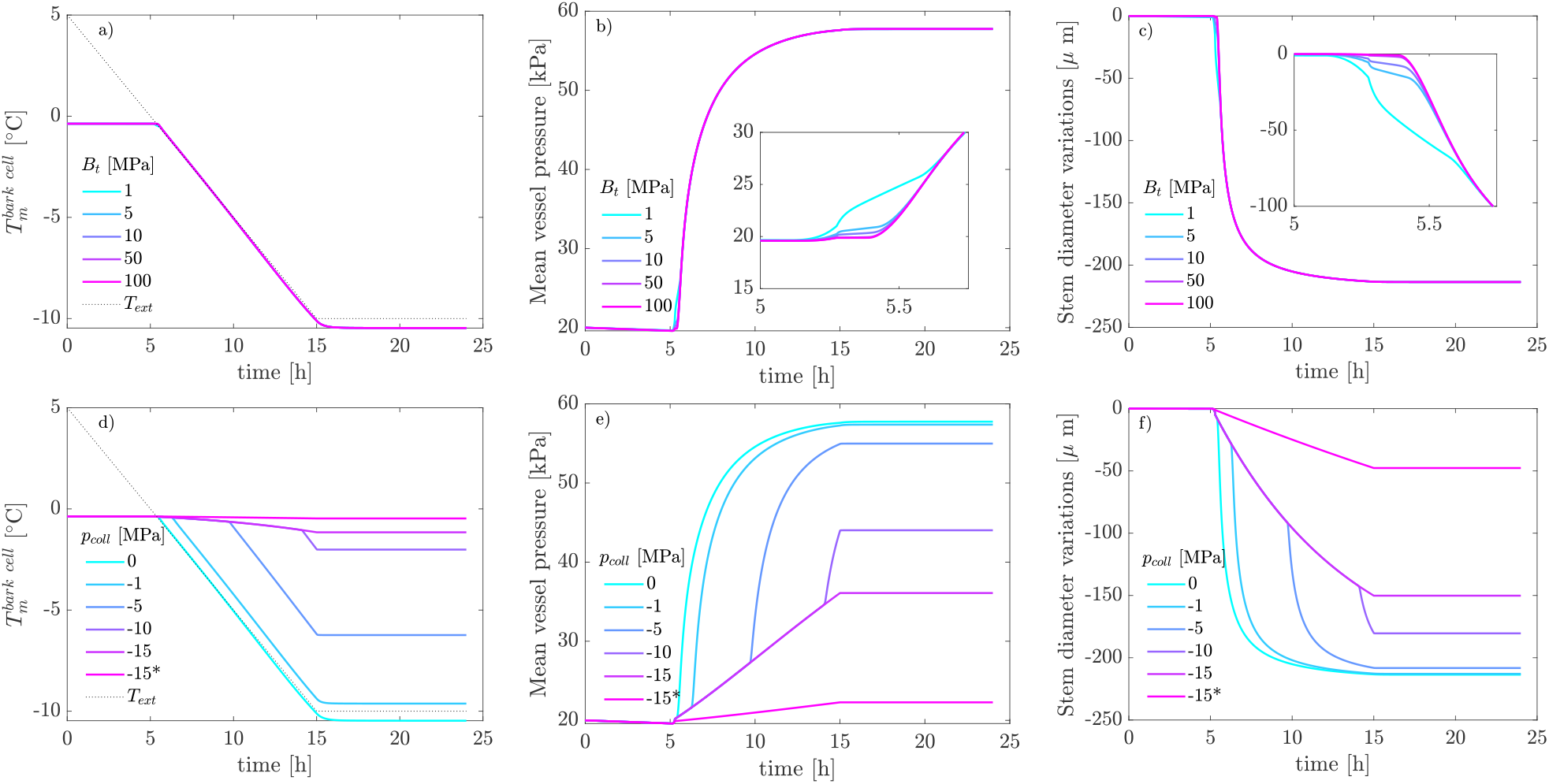
Effect of turgid elastic modulus (a,b,c) and collapse pressure (d,e,f). Unless specified, the parameters are reported in table 2 (base case 2). a-d) bark cell freezing point depression. b-e) Mean vessel water pressure dynamic. c-f) Stem diameter changes. The *\** case in figures d-e-f was computed with *B*_*t*_ = 50 MPa instead of *B*_*t*_ = 10 MPa for the other cases.

In figures 5d-e-f we study the effect of the living cell collaspe pressure (*p*_*coll*_). The decrease in collapse pressure delays turgor loss and allows for the development of negative tugor pressure up to the specified value, at which collapse occurs. As explained in the general presentation of the results, turgor loss point can be seen in figures 5d-e-f as a tipping point between slow and fast dehydration dynamic (and vessel pressure increase). For the lowest collapse pressure, *−*15 MPa, no change can be observed in dehydration speed, meaning that cell collapse, i.e., turgor loss, does not occur. Collapse pressure affects the intracellular freezing avoidance mechanism: the lower the collaspe pressure, the lower the FPD, hence the higher the supercooling degree. For low collapse pressure, an increase in the turgid elastic modulus (*\**case) has now a large effect on FPD, vessel pressure, and stem shrinkage, compared to cases with *p*_*collapse*_ = 0 in figures 5 a-b-c.

### Effect of cooling rate and initial sugar content

The initial living cell sugar content does not affect the final FPD, but its dynamic is significantly different (figure 6a). This is the case for both cooling rate considered here. The initial sugar content does have an effect on the stem shrinkage dynamic, as can be seen in figure 6a: the higher the sugar content, the higher the maximal supercooling degree. For *R*_*c*_ = 1 K/h, the maximal supercooling degree is 0.62°C for the highest initial sugar content (1200 mol/m^3^, 29.1% by mass), compared to 0.08°C for the lowest initial sugar content (200 mol/m^3^, 6.4% by mass). Increasing the cooling rate up to *R*_*c*_ = 8 K/h does also result in an increase of the maximal supercooling degree as it reaches 2.25°C for the highest initial sugar content, and 0.8°C for the lowest initial sugar content.

**Figure 6:**
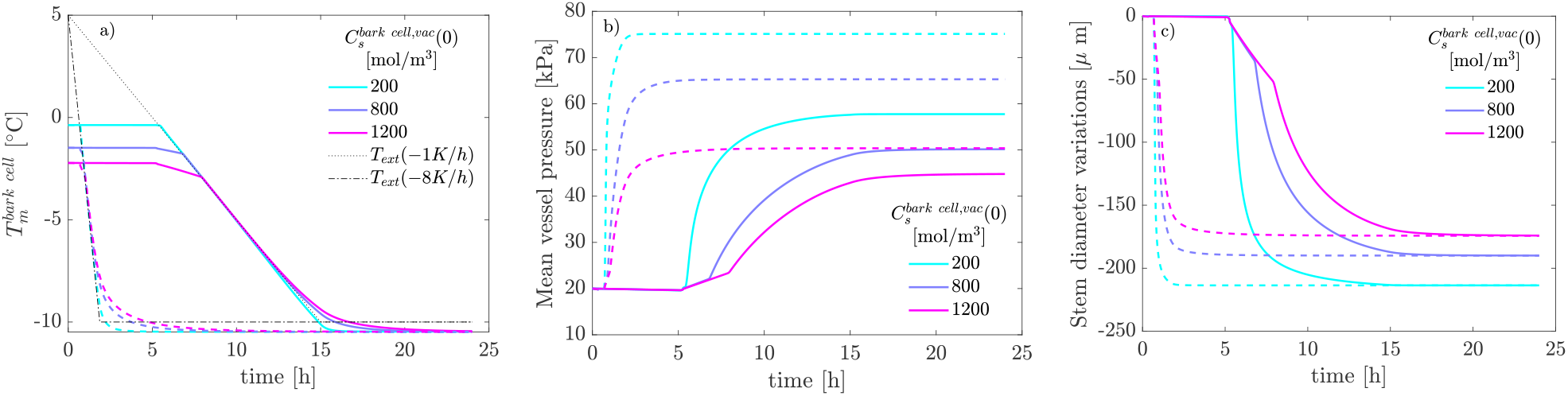
Effect of cooling rate and initial sugar content. Unless otherwise stated, the parameters are reported in table 2 (base case 2). a) Time course of freezing point depression for the bark cell. b) Variation of the average vessel pressure. c) Variation in stem diameter. In a-b-c the solid lines refer to the *R*_*c*_ = 1 K/h case, and the dashed lines to the *R*_*c*_ = 8 K/h case.

Increasing the initial sugar content also results in a decrease of the final mean vessel pressure as well as of a decrease of final stem shrinkage, as seen in figures 6b-c. It also impacts the diameter changes dynamic, with a slower shrinkage for higher initial sugar content (hence a slower FPD dynamic), which also results in a slower pressure increase when increasing sugar content. This is valid for both cooling rate studied here. The cooling rate has no effect on the final diameter shrinkage but does impact the dehydration dynamic, with a much faster shrinkage dynamic for higher cooling rate. On the other hand, the final mean pressure value increases with the cooling rate.

### Effect of external temperature, initial water and sugar content

For all initial water and sugar content, the maximal supercooling, defined as the maximal value of 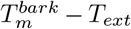, increases with a decrease in *T*_*min*_ (figures 7a-d-g). It also increases with water content, from 2.4°C (*BWF* = 0.0912; 200mol.m^*−*3^; *T*_*min*_ =*−*15°C) up to 13.6°C (*BWF* = 0.2; 200mol.m^*−*3^; *T*_*min*_ =*−*15°C). The effect of initial sugar concentration depends on water fraction value: for the lowest value used in this study, increasing sugar content does slightly increase maximal supercooling degree, similarly to figure 6a, whereas the effect progressively reverses for higher water content. At the highest water content, increasing initial sugar content decreases maximal supercooling degree. Note that the maximal supercooling degree computed here is a transient value and that it can decrease with time depending on *BWF* : for high *BWF* it stabilizes at a value slightly lower than the maximal value and does not decrease towards 0. When increasing *BWF*, the final FPD also becomes sensitive to the initial sugar concentration (the final supercooling degree decreases with an increase in the initial sugar concentration), and not only its dynamic. This is different from what is shown figure 6 for a lower *BWF* (0.0912).

**Figure 7:**
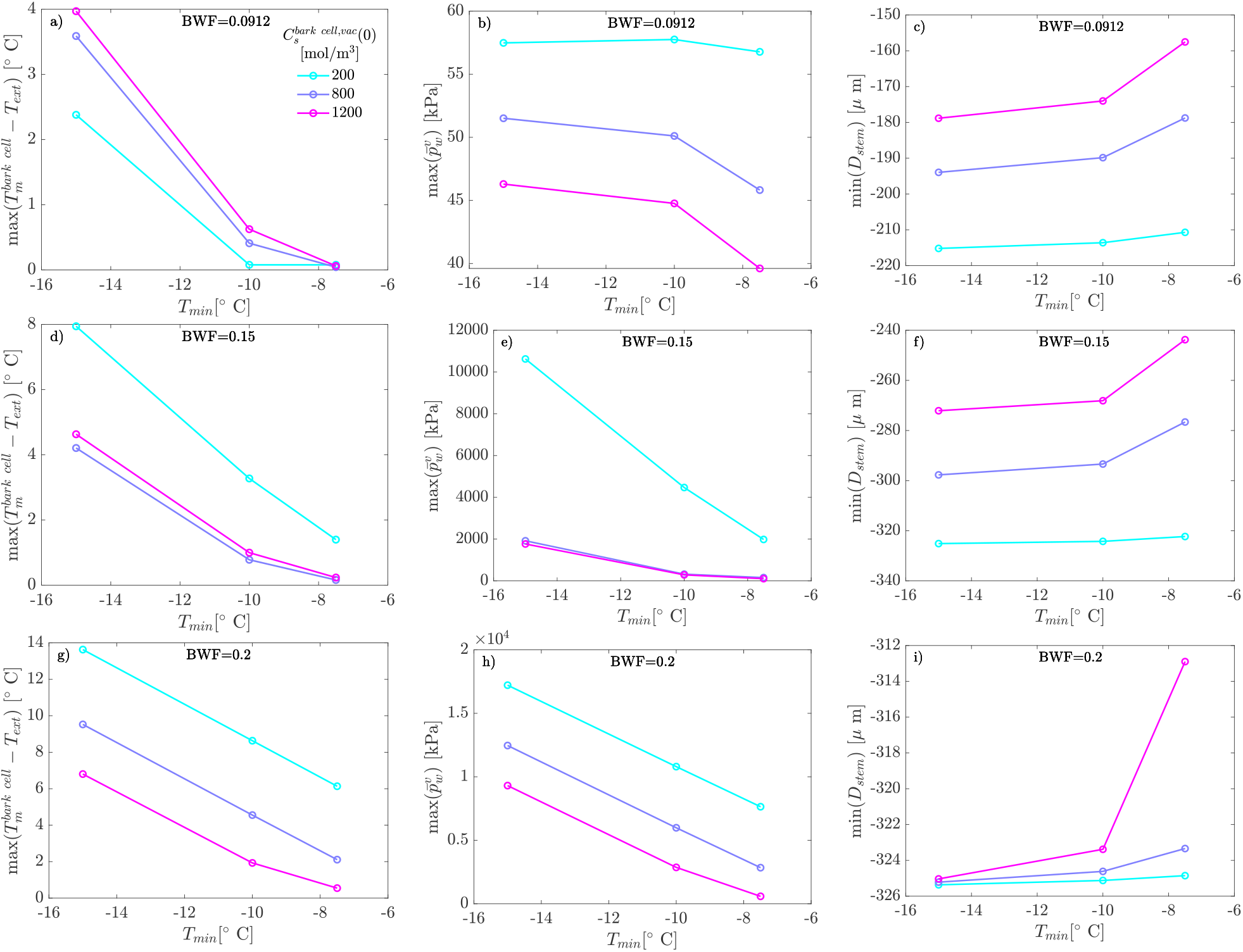
Effect of physiological state (initial living cell sugar content, and bark water fraction) and minimal external temperature on maximal supercooling degree, maximal mean vessel pressure and stem diameter. Unless specified, the parameters are reported in table 2 (base case 2). In each figure the horizontal axis is the minimal external temperature. a-d-g: Maximal bark cell supercooling. b-e-h: Maximal mean vessel water pressure. c-f-i: Minimum stem diameter. BWF value is indicated in each figure, and the colors legend is in 7a.

Similarly, the maximal mean vessel presure, max 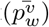, shown in figures 7b-e-h increases with a decrease in *T*_*min*_ for all water and sugar content but one case in figure 7b. It also decreases with sugar content for all water content, and increases with *BWF* over several order of magnitudes. For example, when increasing *BWF* from 0.0912 to 0.2, and at the lowest sugar concentration, it varies from max 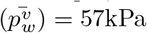to 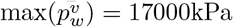. Minimal stem diameter decreases (stem shrinkage increases), with *T*_*min*_, see figure 7c-f-i. Stem shrinkage also increases with *BWF*, and decreases with initial sugar content.

## Discussion

### (Non-)Equilibrium between frozen apoplast and unfrozen symplast

For a living cell to be protected against intracellular freezing by extracellular ice induced dehydration, its phase change temperature must be lower or equal to its temperature, i.e., 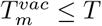 (for a VAC). For *T* = *−*10°C and using Eq. (18), the previous condition leads to a condition for the the VAC osmotic potential: 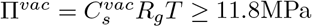. On the other hand, using Eq. (10) for *T* = *−*10°C, the cryostatic suction is 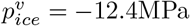. Thus, a sufficient condition for protection against intracelluler freezing becomes

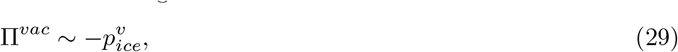

where Π^*vac*^ is the living cell osmotic potential. At equilibrium, apoplast and symplast potentials are equal. Using Eq. (9) and assuming the extracellular ice has zero osmotic potential leads to

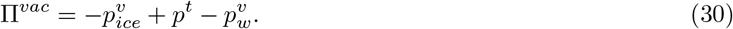

Hence combining the last two equations, we observe that negative *p*^*t*^ (wall tension generation) and positive 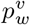can induce 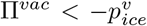. Note that, as shown with the previous example for *T* = *−*10°C, Eq. (29) is often largely sufficient to protect against intracellular freezing, but Eq. (30) is useful to evidence some scenarios that can lead to non-equilibrium freezing, i.e., to the need of supercooling for protection against intracellular freezing. We thus find the following scenarios: (1) if equilibrium is not attained, i.e., if water flow is too slow or impossible, (2) if sufficient cell wall tension is generated, (3) if the apoplast pressure (xylem pressure) is sufficiently high. The model that we developed and validated has been used to explore the effect of environmental, physiological and mechanical parameters that could favour one or several of these different scenarios.

The first scenario, i.e., transient supercooling, is favoured by parameters like cooling rate and initial sugar concentration as seen in figure 6. We limited our study to cooling rates up to 8K/h, which is already high for field conditions. Increasing cooling rate leads to an increased supercooling degree, which is coherent with water flow between symplast and apoplast being too slow to ensure sufficient cell dehydration. The maximal predicted supercooling degree for the highest cooling rate is of 2.25°C, which is low compared to what can be experienced by plant cells (Takahashi et al., 2021). This is coherent with Mazur (1969) who shows that below 10°C/min, plant cells’ survival rate is unaffected by cooling rate. At low water content, an increased initial sugar concentration also leads to an increased supercooling degree (figure 6 and 7a). This might appear as counterintuitive, as increasing sugar content is often described as a hardening mechanism (Poirier et al., 2010; Charrier et al., 2018), but this is coherent with an increased initial viscosity, hence a lower initial hydraulic conductivity, with sugar concentration. Thus, at low water content, our simulations suggest that sugar content is likely to be a resistance mechanism mainly through its effect on nucleation temperature of supercooled cell water rather than through its effect on freeze-induced dehydration.

The second scenario, i.e., steady supercooling due to the generation of cell wall tension has been explored in figure 5. We demonstrated that cell wall elasticity has no effect on dehydration dynamic, xylem pressure nor diameter changes beyond turgor loss point. However, when turgor loss is delayed, cell wall tension may develop before cell collapse occurs, hence giving more importance to cell wall elasticity (figures 5e-f). When cell wall tension develops, dehydration is limited and cell protection against intracellular freezing must rely on supercooling (figure 5d). When cell wall tension is of the order of the cryosuction pressure, almost no dehydration of the cell occurs, as the equilibrium condition, Eq. (30), is ensured without the need for increasing Π^*vac*^. This therefore means that any discussion on the effect of cell wall properties on frost hardiness must be completed by the knowledge of its dehydration extent as done in Stegner et al. (2022). For plant cells avoiding intracellular freezing by dehydration, mechanical properties will have no effects as they quickly lose turgor after freezing inception.

In Stegner et al. (2022), the extent of dehydration was found to correlate negatively with the cell wall elasticity, its thickness and the squared cell wall thickness to cell size ratio. These parameters do indeed influence the collapse pressure of living cells, as demonstrated numerically by Ding et al. (2014); Yang et al. (2017). In the litterature one can find theoretical expressions for collapse pressure that are regularly used for the collapse of vessels/tracheids, but these expressions actually correspond to the threshold pressure for buckling of cylindrical shells under axial compression (Karam, 2005; Jensen and Forterre, 2022). More research is needed to obtain predictive laws for the collapse of cylindrical shells under internal pressure/tension, taking into account their physical properties, dimensions and their complex environment (mechanical cohesion with other cells). It would allow a better prediction of whether supercooling or dehydration is the protection mechanism against intracellular freezing. Such work could be carried out using theoretical and numerical tools.

The third scenario, i.e., steady supercooling due to high vessel pressure has been explored in figure 7. The initial water content is the main parameter driving the increase in vessel pressure: when there is too much ice accumulating in the vessels, the vessel pressure rises up to extremely high values, thus leading to equilibrium, Eq. (30), even for low values of Π^*vac*^. For very high water content, the maximal supercooling degree reaches values up to 14°C. For high water content, increasing sugar content reduces maximal supercooling degree. This is because it gives more weight to the living cell osmotic term compared to the vessel pressure term in Eq. (30). These results on the effect of initial water and sugar content (at high water content) are consistent with frost hardiness results, which has been found to increase with a decrease in water content, and an increase in sugar content (Charrier et al., 2013b, 2018; Baffoin et al., 2021).

Lowering the minimal temperature systematically increases the maximal supercooling degree (figure 7). Whether it is transient or steady depends on cell wall tension or on the water content, as explained previously. But even when it is transient, at the lowest tempererature (*−*15°C) the viscosity is so high, i.e., more than 1000 Pa.s, that the time needed to reach equilibrium can be up to more than 60 hours! In practice, for temperatures lower than *−*10°*C* dehydration is never sufficient to protect cells against intracellular freezing. For such temperature, at the lowest water content, the predicted living cell sugar concentration exceeds 8000 mol/m^3^, which is higher than the supersaturation limit (7152 mol/m^3^, Hartel and Shastry (1991)). Thus, at such low temperatures, sugar precipitation could occur in living cells. However there is, to the best of our knowledge, no evidence that sugar precipitation occurs in plant cells submitted to freezing stress. What could also happen is a glass transition that is known to occur for boreal plants under extremely low temperature (Strimbeck et al., 2015).

### Where is the ice?

Our results show the inter-relation between xylem pressure and stem diameter variations, i.e., high xylem pressure is associated with high stem diameter shrinkage (see figures 3, 5 and 7). This relation is fully expected within the present modelling approach, as water leaving living cells is ending in vessels, where it increases pressure by compressing gas bubbles.

In the model development, we have assumed that ice forms and accumulates in xylem vessels only. This might be a too strong hypothesis as ice could form and accumulate in other places too (xylem fibers or pith, for example). We have also assumed that ice is blocked in vessels until the temperature reaches the phase change temperature. This results in a very sharp stem diameter swelling when the temperature is high enough. On the other hand, in experiments, swelling of the diameter seems more progressive and occurs even at low temperature (Améglio et al., 2001a; Charra-Vaskou et al., 2016; Lindfors et al., 2019). The model predicts that, when vessels are still frozen and the temperature increases, the pressure rises. This is a consequence of the ideal gas law, Eq. (21), and is consistent with the measurements of Améglio et al. (2001b). If ice in the vessels was not blocked and instead allowed to melt under increasing but still negative temperatures, liquid water could back into living cells thus decreasing vessel pressure. It is therefore likely that ice is indeed blocked in the vessels, but the thawing dynamic in other cell types/tissues (fibers, pith) might result in water flow towards the bark at low temperature (below the vessel phase change temperature). Complementary experimental studies are needed to better locate ice in frozen tree stems, at different levels and for different degrees of acclimation.

We hypothesize that ice appears in previously gas filled compartments due to the freezing of a thin water layer covering the walls of these compartments. The resulting ice then draws additional water by cryosuction. This thin water layer could appear due to the condensation of water vapour under decreasing temperature. Water vapour may also condensate at a decreased humidity level in nanopores (Kelvin effect). Ice propagation may also occurs from vessels to intercellular spaces (Ball et al., 2002). We note also that, following Konrad et al. (2019), sublimation may occur, and living cell water could accumulate in the extracellular medium under gaseous form. Under increasing, but still negative, temperatures, the accumulated vapour could condense, or the ice accumulated outside the vessels could melt, allowing liquid water to go back in living cells. These physical processes must be clarified before completing the model.

Turgor loss can be detected in the results of our model as a tipping point in dehydration dynamic: as soon as turgor is lost, living cell dehydration, and thus stem shrinkage, becomes faster (figures 3d-e-f). This tipping point can also be seen in the vessel pressure signals (figures 3a-b). However, in the experiments (figure 4a) no such point was observed. This could result from the fact that cells were not turgid anymore when freezing started. The difference in shrinkage magnitude at short term (figure 4a) might be due to this difference in turgor pressure, but it might more likely be due to the effect of extracellular supercooling. Indeed, in the experiments, freezing occured at a much lower temperature (*−*6°C) than in the model (*−*0.09°C), which resulted in a higher cryosuction at freezing inception, hence a faster shrinkage due to higher flow rate entering vessels.

Reduced stem water content and increased sugar concentration have been found to be linked with the acclimation of a plant to freezing stress (Charrier et al., 2013a,b). In our model, increasing sugar concentration reduces diameter shrinkage, thus reducing vessel pressure (figure 6). This can be understood as follows: with a larger sugar quantity in the cell, the same final sugar concentration is obtained for a larger cell water volume, i.e., as *C*_*s*_ = *n*_*s*_*/V*, the same concentration is obtained for a higher *V* when *n*_*s*_ is higher. Decreasing bark water content leads to reduced stem diameter shrinkage and pressure (figure 7). We have explained in the previous section that decreasing bark water content indeed lead to a better protection by freeze induced dehydration, while increasing sugar content has an effect that depends on water content. The model therefore predicts that non-acclimated trees have a much higher shrinkage than acclimated trees.

On the other hand, in the experiments of Améglio et al. (2001a), non-acclimated trees present smaller stem variations than acclimated trees. Non-acclimated trees present also irreversible stem diameter variations. Such discrepancy could be linked with the effect of acclimation on ice formation location. If ice appears in the bark first, less cell water moves towards the xylem tissue, leading to lower stem shrinkage. Ice could appear in the bark due to intracellular freezing, which would also create irreversible stem shrinkage, or due to ice nucleation in bark intercellular spaces, as seen for *Betula sp* Schott and Roth-Nebelsick (2018).

### Going towards a more accurate phase change modelling

Unlike the method presented in Graf et al. (2015), the present heat transfer method is not a multiscale procedure, where heat transfer and phase changes are described at both the tissue and cellular level. This simplification reduced computational cost and simplified the mathematical description, but does not allow for the computation of fine-scale processes during phase changes, such as volumetric expansion or gas expulsion from the ice crystal. The inclusion of volumetric expansion would induce vessel pressure increase at freezing, and decrease at thawing. In our model, these pressure variations are due to the water flows coming from the living cells. In addition to a better modelling of ice location and accumulation, it would be interesting to apply the method of Graf et al. (2015) to our model, or at least to add an equation to track the ice/water interface (Stefan equation) at the vessel scale, in order to be able to compute the ice volumetric expansion and the associated pressure variations. The inclusion of gas expulsion from the ice/water interface due to low gas solubility in ice could be interesting to draw a link with the generation of gas embolism in xylem vessels (Sevanto et al., 2012). This could for example be done by adding an equation that links the ice/water interface position/speed to the vessel gas quantity, 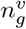.

Finally, the structure of the model offers the possibility to include additional physical processes, e.g., phase separation, eutectic formation, pore effect of FPD (see Esmaeeli et al. (2017) for a numerical model containing such processes), as well as extracellular supercooling (see Meng and Zhang (2020)). Adding extracellular supercooling would additionally allow using the actual thermal conductivity values.

## Conclusion

Our initial aim was to improve the understanding of the competition between freeze-induced dehydration and supercooling for the protection of living cells against intracellular freezing in tree stems. To achieve this, we developed a numerical model that simulates living cell dehydration, xylem pressure and stem diameter changes.

The interests behind this work were multiple. First, the model offered flexibility and control of all parameters, as well as an access to variables that are challenging to measure directly. Second, by explicitly coupling xylem pressure computation and living cell dehydration to stem diameter variations, we facilitated easier comparison of model results with non-intrusive measurements. Third, the impact of any change in the cell dehydration dynamic and extent was observed in pressure and diameter changes, and vice versa. Four, the inclusion of living cell turgor loss in the model allowed us to study the influence of cell wall tension on simulated variables. Finally, as the model includes the impact of physiologically relevant variables, such as water and sugar content, the effect of seasonality and cold acclimation on cell dehydration/supercooling degree, xylem pressure and stem diameter changes was studied.

While adding separately the previously cited missing physiological (more accurate ice location modelling) and physical processes (multiscale phase change, ice/water interface tracking, solution freezing, extracellular supercooling) into the model seems feasible and very interesting, combining them all would pose significant numerical challenges and significantly increase the complexity of the model. Furthermore, the model includes a number of geometrical and anatomical factors that could be used in the future to explore intra and inter-specific differences. In an upcoming work, we intend to employ the model to investigate the mechanisms underlying winter stem pressure build-up in walnut and maple trees.

## Data availability statement

The source code used to generate the data of the present paper can be downloaded at https://github.com/cyrilbz/freezing_branch. Any result of the present paper can be reproduced using this code.

## Acknowledgments

The ANR, through the ACOUFOLLOW project (ANR-20-CE91-0008), is acknowledged for the funding of this project.

## References

G. Alves, J. J. Sauter, J.-L. Julien, P. Fleurat-Lessard, T. Ameglio, A. Guillot, G. Pétel, and A. Lacointe. Plasma membrane H+-ATPase, succinate and isocitrate dehydrogenases activities of vessel-associated cells in walnut trees. Journal of Plant Physiology, 158(10):1263–1271, 2001.

G. Alves, M. Decourteix, P. Fleurat-Lessard, S. Sakr, M. Bonhomme, T. Améglio, A. Lacointe, J.-L. Julien, G. Petel, and A. Guilliot. Spatial activity and expression of plasma membrane H+-ATPase in stem xylem of walnut during dormancy and growth resumption. Tree Physiology, 27(10):1471–1480, 2007.

T. Améglio and P. Cruiziat. Tension-pressure alternation in walnut xylem sap during winter: The role of winter temperature. Comptes Rendus de l’Academie des Sciences Serie 3 Sciences de la Vie (France), 1992.

T. Améglio, H. Cochard, and F. W. Ewers. Stem diameter variations and cold hardiness in walnut trees. Journal of Experimental Botany, 52(364):2135–2142, 2001a.

T. Améglio, F. W. Ewers, H. Cochard, M. Martignac, M. Vandame, C. Bodet, and P. Cruiziat. Winter stem xylem pressure in walnut trees: effects of carbohydrates, cooling and freezing. Tree Physiology, 21(6): 387–394, 2001b.

T. Améglio, C. Bodet, A. Lacointe, and H. Cochard. Winter embolism, mechanisms of xylem hydraulic conductivity recovery and springtime growth patterns in walnut and peach trees. Tree Physiology, 22(17): 1211–1220, 2002.

J. Anderson, L. Gusta, D. Buchanan, and M. Burke. Freezing of water in citrus leaves. Journal of the American Society for Horticultural Science, 108(3):397–400, 1983.

R. Arora. Mechanism of freeze-thaw injury and recovery: A cool retrospective and warming up to new ideas. Plant Science, 270:301–313, 2018.

E. N. Ashworth and F. B. Abeles. Freezing behavior of water in small pores and the possible role in the freezing of plant tissues. Plant Physiology, 76(1):201–204, 1984.

R. Baffoin, G. Charrier, A.-E. Bouchardon, M. Bonhomme, T. Améglio, and A. Lacointe. Seasonal changes in carbohydrates and water content predict dynamics of frost hardiness in various temperate tree species. Tree Physiology, 41(9):1583–1600, 2021.

M. C. Ball, J. Wolfe, M. Canny, M. Hofmann, A. B. Nicotra, and D. Hughes. Space and time dependence of temperature and freezing in evergreen leaves. Functional Plant Biology, 29(11):1259–1272, 2002.

M. C. Ball, M. J. Canny, C. X. Huang, and R. D. Heady. Structural changes in acclimated and unacclimated leaves during freezing and thawing. Functional Plant Biology, 31(1):29–40, 2004.

M. E. Bartolo, S. J. Wallner, and R. E. Ketchum. Comparison of freezing tolerance in cultured plant cells and their respective protoplasts. Cryobiology, 24(1):53–57, 1987.

E. Beck, E.-D. Schulze, M. Senser, and R. Scheibe. Equilibrium freezing of leaf water and extracellular ice formation in afroalpine ‘giant rosette’plants. Planta, 162(3):276–282, 1984.

M. Burke, L. Gusta, H. Quamme, C. Weiser, and P. Li. Freezing and injury in plants. Annual Review of Plant Physiology, 27(1):507–528, 1976.

J. Cavender-Bares. 19 - Impacts of Freezing on Long Distance Transport in Woody Plants. Physiological Ecology. Academic Press, Burlington, 2005. ISBN 978-0-12-088457-5. doi: 10.1016/B978-012088457-5/50021-6. URL https://www.sciencedirect.com/science/article/pii/B9780120884575500216.

M. Ceseri and J. M. Stockie. A mathematical model of sap exudation in maple trees governed by ice melting, gas dissolution, and osmosis. SIAM Journal on Applied Mathematics, 73(2):649–676, 2013.

K. Charra-Vaskou, E. Badel, G. Charrier, A. Ponomarenko, M. Bonhomme, L. Foucat, S. Mayr, and T. Améglio. Cavitation and water fluxes driven by ice water potential in Juglans regia during freeze–thaw cycles. Journal of Experimental Botany, 67(3):739–750, 2016.

G. Charrier, H. Cochard, and T. Améglio. Evaluation of the impact of frost resistances on potential altitudinal limit of trees. Tree Physiology, 33(9):891–902, 09 2013a.

G. Charrier, M. Poirier, M. Bonhomme, A. Lacointe, and T. Améglio. Frost hardiness in walnut trees (Juglans regia L.): how to link physiology and modelling? Tree Physiology, 33(11):1229–1241, 2013b.

G. Charrier, J. Ngao, M. Saudreau, and T. Améglio. Effects of environmental factors and management practices on microclimate, winter physiology, and frost resistance in trees. Frontiers in Plant Science, 6:259, 2015.

G. Charrier, A. Lacointe, and T. Améglio. Dynamic modeling of carbon metabolism during the dormant period accurately predicts the changes in frost hardiness in walnut trees juglans regia l. Frontiers in Plant Science, 9:1746, 2018.

F. Chenlo, R. Moreira, G. Pereira, and A. Ampudia. Viscosities of aqueous solutions of sucrose and sodium chloride of interest in osmotic dehydration processes. Journal of Food Engineering, 54(4):347–352, 2002.

H. Cochard, N. Bréda, A. Granier, and G. Aussenac. Vulnerability to air embolism of three european oak species (Quercus petraea (matt) liebl, Q pubescens willd, Q robur l). In Annales des Sciences Forestières, volume 49, pages 225–233. EDP Sciences, 1992.

H. Cochard, D. Lemoine, T. Améglio, and A. Granier. Mechanisms of xylem recovery from winter embolism in Fagus sylvatica. Tree Physiology, 21(1):27–33, 2001.

COMSOL. version 5.6, 2020. URL http://www.comsol.com/products/multiphysics/.

Y. Ding, Y. Zhang, Q.-S. Zheng, and M. T. Tyree. Pressure–volume curves: revisiting the impact of negative turgor during cell collapse by literature review and simulations of cell micromechanics. New Phytologist, 203(2):378–387, 2014.

J. Dumais and Y. Forterre. “vegetable dynamicks”: the role of water in plant movements. Annual Review of Fluid Mechanics, 44:453–478, 2012.

H. S. Esmaeeli, Y. Farnam, D. P. Bentz, P. D. Zavattieri, and W. J. Weiss. Numerical simulation of the freeze–thaw behavior of mortar containing deicing salt solution. Materials and Structures, 50:1–20, 2017.

L. Eurich, R. Schott, S. Shahmoradi, A. Wagner, R. I. Borja, A. Roth-Nebelsick, and W. Ehlers. A thermo-dynamically consistent quasi-double-porosity thermo-hydro-mechanical model for cell dehydration of plant tissues at subzero temperatures. Archive of Applied Mechanics, pages 1–29, 2021.

F. W. Ewers, T. Ameglio, H. Cochard, F. Beaujard, M. Martignac, M. Vandame, C. Bodet, and P. Cruiziat. Seasonal variation in xylem pressure of walnut trees: root and stem pressures. Tree Physiology, 21(15): 1123–1132, 2001.

I. Graf, M. Ceseri, and J. M. Stockie. Multiscale model of a freeze–thaw process for tree sap exudation. Journal of the Royal Society Interface, 12(111):20150665, 2015.

M. Griffith and G. N. Brown. Cell wall deposits in winter rye Secale cereale L.’puma’during cold acclimation. Botanical Gazette, 143(4):486–490, 1982.

M. Griffith and M. W. Yaish. Antifreeze proteins in overwintering plants: a tale of two activities. Trends in Plant Science, 9(8):399–405, 2004.

R. W. Hartel and A. V. Shastry. Sugar crystallization in food products. Critical Reviews in Food Science & Nutrition, 30(1):49–112, 1991.

T. Hölttä, T. Vesala, M. Perämäki, and E. Nikinmaa. Refilling of embolised conduits as a consequence of ‘Münch water’circulation. Functional Plant Biology, 33(10):949–959, 2006.

M. Ishikawa and A. Sakai. Extraorgan freezing in wintering flower buds of Cornus officinalis Sieb. et Zucc. Plant, Cell & Environment, 8(5):333–338, 1985.

K. Jensen and Y. Forterre. Soft Matter in Plants: From Biophysics to Biomimetics, volume 15. Royal Society of Chemistry, 2022.

G. N. Karam. Biomechanical model of the xylem vessels in vascular plants. Annals of Botany, 95(7):1179–1186, 2005.

W. Konrad, R. Schott, and A. Roth-Nebelsick. A model for extracellular freezing based on observations on Equisetum hyemale. Journal of Theoretical Biology, 478:161–168, 2019.

I. Leinonen. A simulation model for the annual frost hardiness and freeze damage of scots pine. Annals of Botany, 78(6):687–693, 1996.

J. Levitt et al. Responses of Plants to Environmental Stress, Volume 1: Chilling, Freezing, and High Temperature Stresses. Academic Press., 1980.

L. Lindfors, J. Atherton, A. Riikonen, and T. Hölttä. A mechanistic model of winter stem diameter dynamics reveals the time constant of diameter changes and the elastic modulus across tissues and species. Agricultural and Forest Meteorology, 272:20–29, 2019.

A. Lintunen, L. Lindfors, E. Nikinmaa, and T. Hölttä. Xylem diameter changes during osmotic stress, desiccation and freezing in Pinus sylvestris and Populus tremula. Tree Physiology, 37(4):491–500, 2017.

J. Loch. Thermodynamic equilibrium between ice and water in porous media. Soil Science, 126(2):77–80, 1978.

MATLAB. version 9.4.0 (R2018b). The MathWorks Inc., Natick, Massachusetts, 2018.

P. Mazur. Freezing injury in plants. Annual Review of Plant Physiology, 20(1):419–448, 1969.

Z. Meng and P. Zhang. Dynamic propagation of ice-water phase front in a supercooled water droplet. International Journal of Heat and Mass Transfer, 152:119468, 2020.

J. Milburn and P. O’Malley. Freeze-induced sap absorption in Acer pseudoplatanus: a possible mechanism. Canadian Journal of Botany, 62(10):2101–2106, 1984.

G. Neuner, B. Xu, and J. Hacker. Velocity and pattern of ice propagation and deep supercooling in woody stems of Castanea sativa, Morus nigra and Quercus robur measured by IDTA. Tree Physiology, 30(8):1037–1045, 2010.

G. Neuner, K. Monitzer, D. Kaplenig, and J. Ingruber. Frost survival mechanism of vegetative buds in temperate trees: deep supercooling and extraorgan freezing vs. ice tolerance. Frontiers in Plant Science, 10:537, 2019.

C. R. Olien. Freezing stresses and survival. Annual Review of Plant Physiology, 18(1):387–408, 1967.

P. O’Malley and J. Milburn. Freeze-induced fluctuations in xylem sap pressure in Acer pseudoplatanus. Canadian Journal of Botany, 61(12):3100–3106, 1983.

R. S. Pearce. Plant freezing and damage. Annals of Botany, 87(4):417–424, 2001.

J. Petty and M. A. Palin. Permeability to water of the fibre cell wall material of two hardwoods. Journal of Experimental Botany, 34(6):688–693, 1983.

M. Poirier, A. Lacointe, and T. Ameglio. A semi-physiological model of cold hardening and dehardening in walnut stem. Tree Physiology, 30(12):1555–1569, 2010.

R. Prapainop and K. Maneeratana. Simulation of ice formation by the finite volume method. Simulation, 26(1): 56, 2004.

C. Rajashekar and M. J. Burke. Freezing characteristics of rigid plant tissues (development of cell tension during extracellular freezing). Plant Physiology, 111(2):597–603, 1996.

D. Robson and J. Petty. Freezing in conifer xylem: I. pressure changes and growth velocity of ice. Journal of Experimental Botany, 38(11):1901–1908, 1987.

A. Sakai and W. Larcher. Frost survival of plants: responses and adaptation to freezing stress, volume 62. Springer Science & Business Media, 2012.

R. T. Schott and A. Roth-Nebelsick. Ice nucleation in stems of trees and shrubs with different frost resistance. IAWA Journal, 39(2):177–190, 2018.

R. T. Schott, D. Voigt, and A. Roth-Nebelsick. Extracellular ice management in the frost hardy horsetail Equisetum hyemale L. Flora, 234:207–214, 2017.

S. Sevanto, N. M. Holbrook, and M. C. Ball. Freeze/thaw-induced embolism: probability of critical bubble formation depends on speed of ice formation. Frontiers in Plant Science, 3:107, 2012.

D. Siminovitch and G. Scarth. A study of the mechanism of frost injury to plants. Canadian Journal of Research, 16(11):467–481, 1938.

M. Stefanowska, M. Kuraś, and A. Kacperska. Low temperature-induced modifications in cell ultrastructure and localization of phenolics in winter oilseed rape (Brassica napus L. var. oleifera L.) leaves. Annals of Botany, 90(5):637–645, 2002.

M. Stegner, A. Flörl, J. Lindner, S. Plangger, T. Schaefernolte, A.-L. Strasser, V. Thoma, J. Walde, and G. Neuner. Freeze dehydration vs. supercooling of mesophyll cells: Impact of cell wall, cellular and tissue traits on the extent of water displacement. Physiologia Plantarum, 174(6):e13793, 2022.

E. Steudle, U. Zimmermann, and U. Lüttge. Effect of turgor pressure and cell size on the wall elasticity of plant cells. Plant Physiology, 59(2):285–289, 1977.

G. R. Strimbeck, P. G. Schaberg, C. G. Fossdal, W. P. Schröder, and T. D. Kjellsen. Extreme low temperature tolerance in woody plants. Frontiers in Plant Science, 6:884, 2015.

D. Takahashi, I. R. Willick, J. Kasuga, and D. P. Livingston III. Responses of the plant cell wall to sub-zero temperatures: a brief update. Plant and Cell Physiology, 62(12):1858–1866, 2021.

M. Toner, E. G. Cravalho, and M. Karel. Thermodynamics and kinetics of intracellular ice formation during freezing of biological cells. Journal of Applied Physics, 67(3):1582–1593, 1990.

L. E. Towill and P. Mazur. Osmotic shrinkage as a factor in freezing injury in plant tissue cultures. Plant Physiology, 57(2):290–296, 1976.

M. T. Tyree. Maple sap uptake, exudation, and pressure changes correlated with freezing exotherms and thawing endotherms. Plant Physiology, 73(2):277–285, 1983.

Y. Utsumi, Y. Sano, S. Fujikawa, R. Funada, and J. Ohtani. Visualization of cavitated vessels in winter and refilled vessels in spring in diffuse-porous trees by cryo-scanning electron microscopy. Plant Physiology, 117 (4):1463–1471, 1998.

M. Wisniewski. Deep supercooling in woody plants and the role of cell wall structure. Biological ice nucleation and its applications, pages 163–181, 1995.

D. Yang, S. Pan, Y. Ding, and M. T. Tyree. Experimental evidence for negative turgor pressure in small leaf cells of Robinia pseudoacacia L versus large cells of Metasequoia glyptostroboides Hu et WC Cheng. 1. evidence from pressure-volume curve analysis of dead tissue. Plant, Cell & Environment, 40(3):351–363, 2017.

R. Zweifel and R. Häsler. Frost-induced reversible shrinkage of bark of mature subalpine conifers. Agricultural and Forest Meteorology, 102(4):213–222, 2000.

